# Increasing the compositional heterogeneity of single-chain amphiphile membranes supported by coacervate cores alters stability and properties of the hybrid protocells

**DOI:** 10.64898/2026.02.02.703427

**Authors:** Manesh Prakash Joshi, Jessica Lee, Maxfield Chan, Christine D. Keating

**Affiliations:** Department of Chemistry, The Pennsylvania State University, University Park, PA 16802 USA

## Abstract

Coacervate droplets and lipid vesicles are two classes of self-assembled compartments that have been proposed as protocell models. Hybrid protocells, in which a coacervate core is surrounded by a lipid membrane, can integrate the advantages of both protocell systems while overcoming their limitations. Although hybrid protocell membranes have been produced with a variety of diacyl phospholipids related to modern biology and some single-chain amphiphiles inspired by prebiotic scenarios, little is known about how mixtures of single-chain amphiphiles impact hybrid protocell membrane formation and properties. Given the plausible diversity of amphiphiles in the prebiotic milieu, the resulting membranes would have inherently incorporated multiple lipids of different types, potentially altering the properties and viability of hybrid protocells in their environment. Here, we systematically increased the compositional heterogeneity of hybrid protocell membranes by using different prebiotically relevant single-chain amphiphiles of varying head groups and alkyl chain lengths. These membranes were assembled around model coacervate droplets generated from polyallylamine hydrochloride and adenosine diphosphate, and the effect of heterogeneity on membrane properties and stability was evaluated. Compared to protocells with homogeneous membranes, those with heterogeneous amphiphile membranes exhibited higher yields, smaller sizes, and greater sub-compartment formation. Also, they showed increased membrane order, retained similar lateral lipid diffusion, and showed population-level variability in permeability to small anionic molecules. Notably, heterogeneous membranes showed enhanced structural stability under acidic conditions, retaining key properties like size and sub-compartment heterogeneity, thereby broadening the pH range over which hybrid protocells remain intact. These findings suggest that amphiphile diversity not only would have influenced the structural properties of hybrid protocells but also created diversity within the protocell population and enhanced their robustness, thereby playing a crucial role in protocell evolution on early Earth.

## INTRODUCTION

Protocells are primitive, cell-like structures hypothesized to bridge the gap between non-living matter and living systems. They are central to studies on the origin of life, offering insights into how cellular life could emerge on Earth or other Earth-like planets.^1–3^ In this context, vesicles and coacervates are widely studied as model protocell systems.^1,4^ Vesicles are generated by the spontaneous self-assembly of prebiotically plausible single-chain amphiphiles (SCAs) like fatty acids.^5,6^ They provide stable, semipermeable boundary conditions, protecting the luminal content from the surrounding medium, similar to modern cell membranes.^7,8^ However, they lack the molecular crowding typical of cytoplasm and generally exhibit low encapsulation efficiency.^9^ Coacervates form through the associative liquid-liquid phase separation of macromolecules or other “sticky” solutes that can include oligopeptides, nucleotides, or even SCAs.^9–14^ The resulting liquid droplets provide a molecularly crowded proto-cytoplasmic environment, and locally concentrate molecules sequestered from the larger bulk solution. However, coacervate droplets are intrinsically unstable with respect to coalescence, and in many cases, molecular cargo can transfer between droplets by diffusion, resulting in loss of their individual protocell identity over time.^15–17^

To combine the advantageous features of vesicles and coacervates while overcoming their limitations, Tang et al. proposed a new model protocell system, the hybrid protocell (HP).^18^ It is a structural amalgamation of vesicles and coacervates, where a microscale coacervate droplet is surrounded by a lipid membrane layer. This outer membrane provides a semipermeable barrier that protects and stabilizes the inner coacervate core by preventing its coalescence, whereas the coacervate creates a molecularly crowded environment that efficiently encapsulates molecules during HP formation. Due to their useful features, hybrid protocells (HPs) have garnered growing interest in fields beyond the origin of life research, including pharmaceutical sciences, synthetic biology, soft matter, and bioinspired material chemistry.^19–26^ Most studies on HPs have used diacyl phospholipids to generate HP membranes.^20,23,26^ In the context of the origin of life research, HP membranes have been generated using fatty acids because of their prebiotic relevance and ability to self-assemble into membrane bilayers.^5,18,27^

However, fatty acids were likely not the only amphiphiles present on early Earth. Experimental evidence^28,29^ suggests that a “prebiotic soup” would have contained a diverse set of amphiphiles, in addition to fatty acids, with distinct physicochemical properties. Consequently, prebiotic membranes generated from these amphiphiles were also likely compositionally heterogeneous. Importantly, this heterogeneity may have played a critical role in shaping protocell properties, including their formation, stability, and function. Several studies on vesicle-based protocells (membranes without a coacervate core) have highlighted the significance of membrane heterogeneity. For example, increasing compositional diversity can reduce the critical vesicle concentration (CVC) of the amphiphile system,^30,31^ and also facilitate the membrane assembly under natural, early Earth analog hot spring conditions.^32^ It can also increase the pH, temperature, and metal ion stability of protocell membranes^31,33–35^ and modulate their physicochemical properties like permeability and membrane order.^36,37^ Furthermore, protocells with compositionally heterogeneous membranes have been shown to extract lipids from those with homogeneous membranes, enabling them to undergo growth and division,^38^ mimicking an important feature of living systems.

Although membrane heterogeneity has been extensively investigated using vesicles as a model protocell system (as discussed above), its influence on more complex and advanced systems such as hybrid protocells remains largely unexplored. Recently, we demonstrated how the presence of a coacervate core can remarkably affect the properties of surrounding membranes made of either pure fatty acids or fatty acid/phospholipid blended systems.^27^ These coacervate-supported membranes showed significantly different permeability behavior and enhanced stability in the presence of Mg^2+^ compared to the corresponding vesicles, likely due to charge-charge interactions between anionic carboxylate groups of the fatty acids and cationic amine moieties of the coacervates. We were interested to learn whether hybrid protocells could accommodate compositionally heterogeneous membranes assembled from prebiotically relevant SCAs, and if so, how such heterogeneity influenced their physicochemical properties, stability, and potential functional capabilities.

Here, we describe the formation of HPs having SCA membranes of increasing compositional heterogeneity, both in terms of the amphiphile head group and chain length, and compare the properties of homogeneous and heterogeneous HPs. Specifically, the influence of heterogeneity on HP yield and key physicochemical properties, including size, sub-compartment formation, membrane order, lateral diffusion, and membrane permeability, was evaluated. Heterogeneous HPs showed overall higher yield, smaller size, and enhanced sub-compartment formation efficiency than homogeneous HPs. Also, their membranes exhibited increased order and variable permeability within the population toward small anionic molecules, such as fluorescein. We also examined how membrane heterogeneity affects the stability of HPs under fluctuating pH conditions mimicking those of early Earth hydrothermal systems. While both homogeneous and heterogeneous HPs remained stable in alkaline environments, the latter showed enhanced stability under acidic conditions, while retaining the key structural features, such as size and sub-compartment heterogeneity. Overall, our study demonstrates a more realistic model of HPs by incorporating prebiotic amphiphile diversity and complexity and provides new insight into how membrane heterogeneity could have shaped the early stages of protocell evolution.

## RESULTS AND DISCUSSION

### Generation of hybrid protocells with a compositionally heterogeneous membrane

Firstly, we tested whether HPs with compositionally heterogeneous membranes can be generated from a mixture of simple SCAs. The SCAs used here included oleic acid (OA), glycerol-1-monooleate (GMO), and oleyl alcohol (OOH) (Figure 1B), which are frequently used to generate model protocell membranes without a coacervate core.^31,32,39,40^ Different combinations of these amphiphiles, including OA (9 mM), OA + GMO (9 mM; 2:1 ratio), and OA + GMO + OOH (9 mM; 4:1:1 ratio), were tested for HP formation. Notably, all these combinations are known to generate vesicles.^31,32^ We note that OA, but not GMO or OOH, can self-assemble into membrane bilayers on its own under the experimental conditions we used.^41–43^ An amphiphile system containing only OA would generate HPs with a homogeneous membrane (made of OA). Contrarily, the other two amphiphile systems (OA + GMO and OA + GMO + OOH) would generate HPs with compositionally heterogeneous membranes containing a mixture of fatty acids and other amphiphiles (Figure 1A). The membranes were assembled around the surface of a coacervate core made of a structurally simple model polycation (polyallylamine hydrochloride, PAH) and prebiotically relevant adenosine diphosphate (ADP). PAH and ADP (40 mM total charge; 1:1 charge ratio) (Figure 1B) were used as a model system to generate the coacervate component of HPs since PAH/ADP coacervates are known to support membrane formation around their surface.^27^ The HPs were prepared by a gentle hydration method where a lipid film was hydrated with a PAH/ADP coacervate solution prepared in 100 mM bicine buffer pH 8.5. The membrane and the coacervate parts of HPs were stained with Octadecyl Rhodamine B Chloride (R18) and Alexa Fluor 488-labeled PAH, respectively, to differentially visualize these components under the fluorescence microscope.

**Figure 1.**
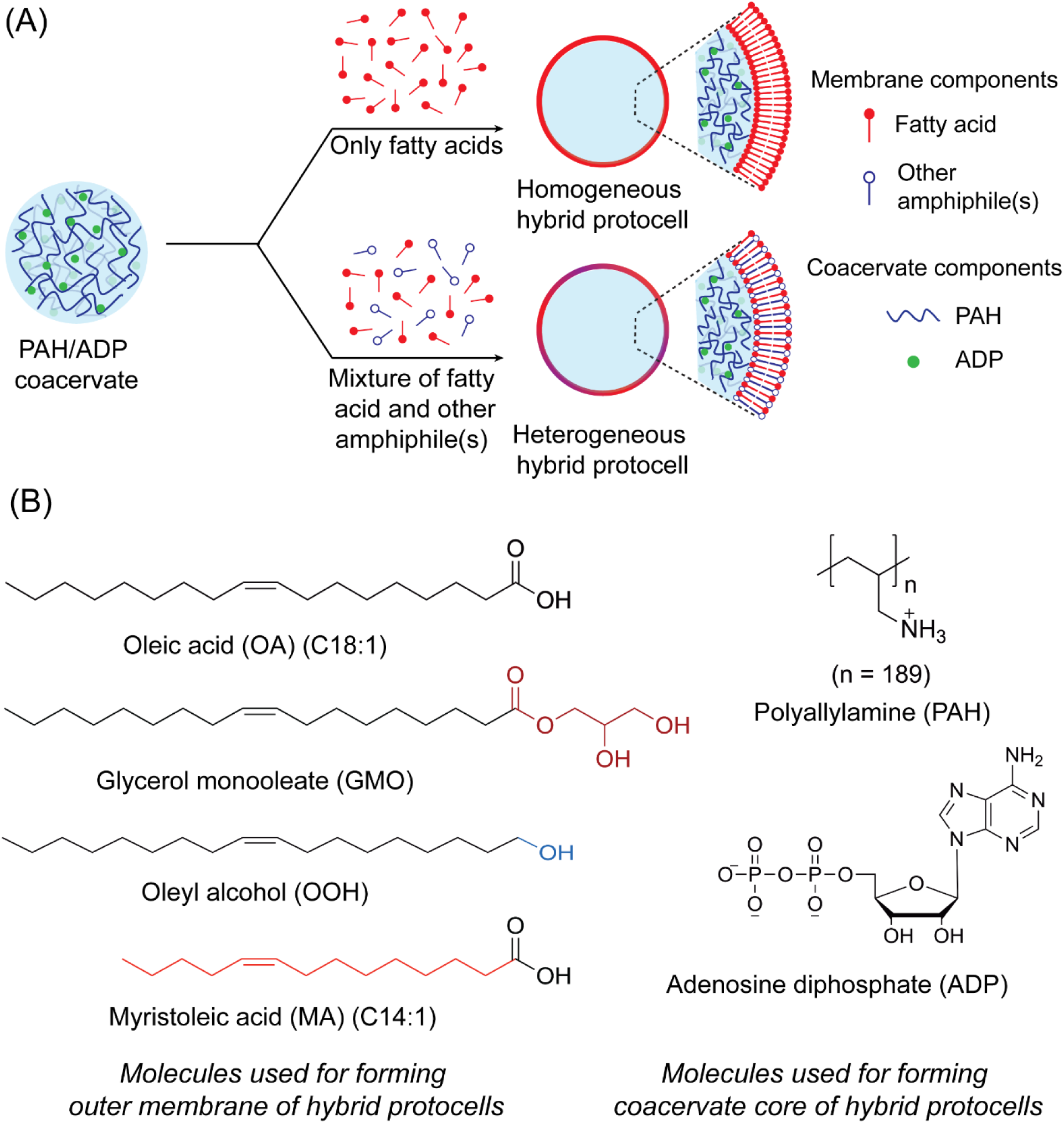
Overview of hybrid protocell formation and the molecules used in this study. **(A)** Schematic illustration of homogeneous and heterogeneous HPs formed by coating PAH/ADP coacervates with single-chain amphiphile membranes. **(B)** Chemical structures of different molecules used for generating HPs. Differences in head group and alkyl chain length of amphiphiles relative to oleic acid are highlighted in different colors (left panel).

All three amphiphile systems mentioned above were able to generate HPs where a PAH/ADP coacervate core was surrounded by a fatty acid-based membrane (Figure 2A). This was evident by the localization of membrane-staining R18 dye around the coacervate surface. Furthermore, the Alexa Fluor 488-labeled PAH dye was localized in the lumen, confirming the coacervate encapsulation by the membrane. Also, the coacervate dye was found to preferentially accumulate on the membrane-coacervate interface (see white arrows in the coacervate panel of Figure 2A). We note that this accumulation was not due to its interaction with the membrane dye (Figure S1) but was likely due to electrostatic interactions with negatively charged fatty acids in the membrane. In addition to HPs, a few vesicles lacking a coacervate core were also present in the solution. Interestingly, we did not observe any free PAH/ADP coacervates (without membranes), indicating that a 9 mM total amphiphile concentration was sufficient to generate membranes that coated all available coacervate surfaces. However, at lower total amphiphile concentration, such as 3 mM OA, the amphiphiles were predominantly sequestered within the coacervates rather than forming a membrane around them (Figure S2). Consequently, very few HPs were observed at this amphiphile concentration. Such sequestration of amphiphiles into coacervates at low amphiphile concentrations has been reported previously.^44^ Notably, 3 mM OA is well above its critical vesicle (bilayer) concentration, which lies in the range of ∼0.2–0.7 mM at pH 8.5 under low-salt conditions.^31,45^ This suggests that the presence of coacervates substantially alters the membrane self-assembly behavior of SCAs, such that higher amphiphile concentrations are required to form HP membranes than to generate free fatty-acid vesicles.

**Figure 2.**
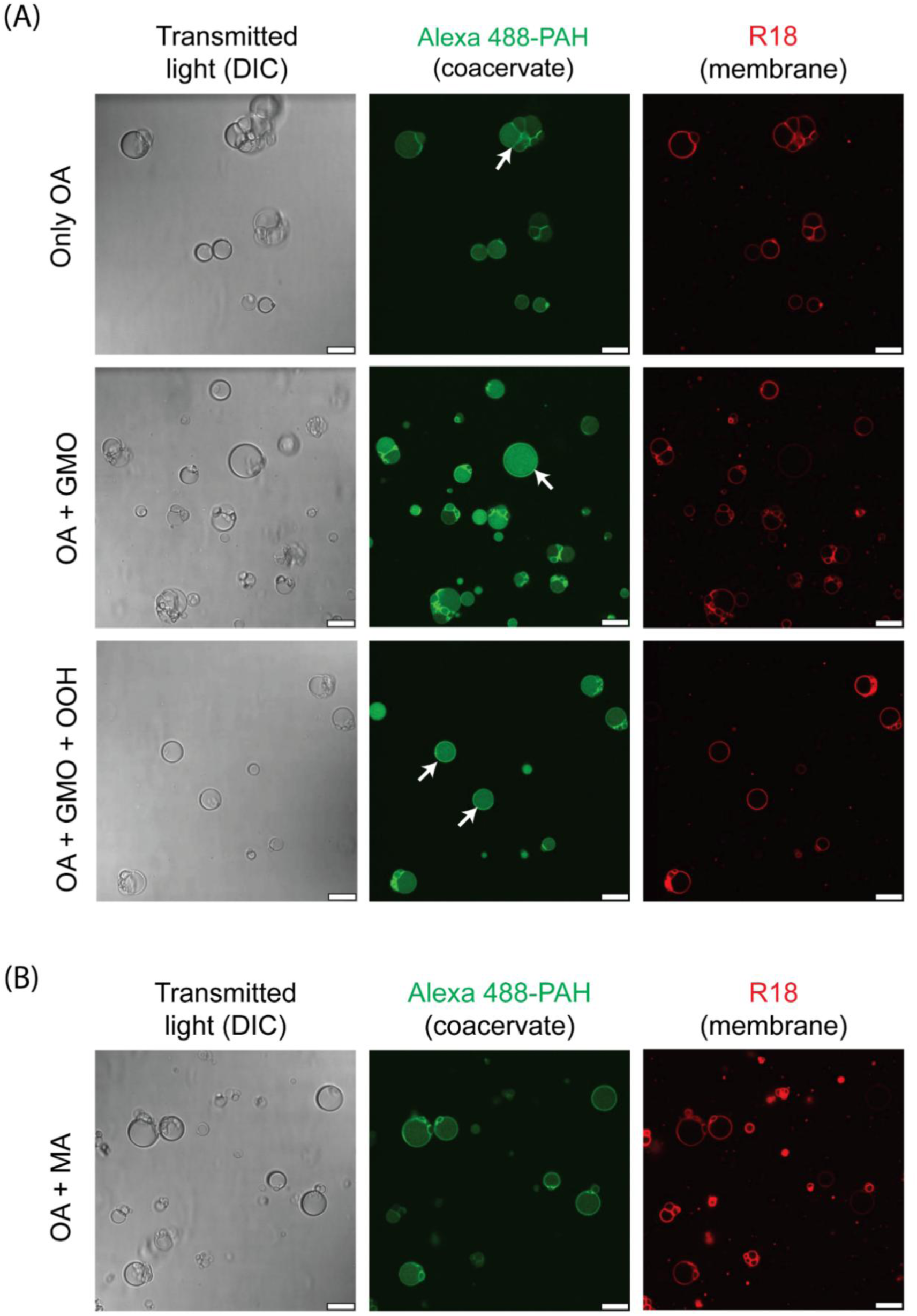
Formation of hybrid protocells with compositionally heterogeneous membranes. **(A)** Confocal microscopy images showing the formation of HPs with homogeneous and heterogeneous membrane compositions. Homogeneous membranes are made of 9 mM OA, whereas heterogeneous membranes are composed of different SCA combinations having variability in the head group, such as OA + GMO (9 mM, 2:1 ratio) and OA + GMO + OOH (9 mM; 4:1:1 ratio). **(B)** Formation of HPs by the mixture of OA and MA (15 mM; 1:9 ratio), having variability in the alkyl chain length (OA (C18:1) and MA (C14:1)). Both types of membranes are self-assembled around PAH/ADP coacervates (40 mM total charge; 1:1 charge ratio) prepared in 100 mM bicine buffer pH 8.5. Differential Interference Contrast (DIC, left panels) images show the actual morphology of HPs without staining, whereas the membrane and coacervate parts of the HP are visualized by fluorescent labeling with R18 (red color, right panels) and Alexa Fluor 488-PAH (green color, middle panels), respectively. White arrows indicate HPs showing the preferential localization of a coacervate dye on the membrane-coacervate interface. Fluorescence images have been pseudo-colored and contrast-adjusted for better visualization. All scale bars are 10 µm.

Overall, these results demonstrated that HPs with compositionally heterogeneous membranes could be generated from simple SCAs that are used to form model protocell membranes. Our previous work showed that PAH/ADP coacervates possess a net positive surface charge, which promotes the assembly of negatively charged fatty acid/phospholipid membranes via electrostatic interactions.^27^ A similar mechanism likely underlies membrane formation around a coacervate here. In general, the incorporation of uncharged lipids such as GMO and OOH into OA membranes reduces the overall surface negative charge density of the membrane,^31^ which could potentially diminish its capacity to interact with coacervate polycations. Nevertheless, these mixed SCA systems were still able to form membranes around the coacervates, showing the robustness of this assembly process to changes in lipid composition.

Amphiphiles typically consist of two distinct structural regions: a polar (hydrophilic) head group and a non-polar (hydrophobic) aliphatic tail. The HPs generated earlier had heterogeneity in the amphiphile head group region in terms of the size and the polarity of the head group (Figure 1B). Such heterogeneity could also be observed in terms of the hydrophobic tail of amphiphiles in a mixture of fatty acids with different chain lengths. This chain-length heterogeneity also seems rational from a prebiotic perspective since the analysis of the plausible exogenous^28^ and endogenous^29^ sources of amphiphiles on early Earth showed the presence of fatty acids with varying chain lengths.

Therefore, we tested whether the HPs with membrane heterogeneity in the alkyl chain length of membrane lipids could be generated from a mixture of different chain-length fatty acids. For this, a lipid film containing oleic acid (C18:1) and myristoleic acid (MA; C14:1) (15 mM; 1:9 ratio of OA to MA) (structures shown in Figure 1B) was hydrated with 100 mM tris buffer, pH 8. The ratio of OA to MA and the solution pH were selected based on the previous studies on the vesicle formation behavior of this mixed amphiphile system.^46^ Similar to head-group heterogeneity experiments, the mixture of OA and MA was also able to form HPs with PAH/ADP coacervates, with a very good yield (Figure 2B), indicating that the heterogeneity can also be introduced in the amphiphile chain length.

So far, we demonstrated that HPs with compositionally heterogeneous membranes can be generated from a mixture of SCAs with varying head groups and alkyl chain lengths. However, the microscopic analysis used to confirm HP formation does not provide any information about the heterogeneous nature of the membrane, as both the homogeneous and the heterogeneous HP membranes appear similar under the microscope (Figures 2 and S3). This could lead to a false-positive result regarding membrane heterogeneity, since OA itself can independently generate an HP membrane. Therefore, we confirmed the membrane heterogeneity of HPs by specifically detecting the presence of amphiphiles in the HP membrane, after separating them from other potential amphiphile sources (vesicles and free amphiphiles) in the solution. To achieve this, we developed a protocol in which HPs were separated from vesicles and free amphiphiles by centrifugation, where the HPs, containing a dense coacervate core, settled as a pellet, while vesicles and free amphiphiles remained in the supernatant (Figures S4 and S5). Notably, this pellet was not observed in the control reaction containing only vesicles (Figure S5). Subsequently, the HP pellet was resuspended in the same buffer, which was used for their preparation, and amphiphiles present in the HP membrane were separated from coacervate-forming molecules by butanol extraction, where amphiphiles preferentially go into the butanol phase. The presence of amphiphiles in the butanol phase was confirmed by thin-layer chromatography (TLC) and mass spectrometry.

First, we confirmed the heterogeneity of OA + GMO HPs using the above-mentioned protocol. The preliminary TLC analysis of the butanol phase after extraction showed two distinct spots, which were comparable to those of the OA and GMO standards (Figure 3A), indicating the presence of both OA and GMO in the HP membrane. This was further confirmed by mass analysis, where the expected masses corresponding to both OA (281.2480) and GMO (401.2898) (Figure 3C) were detected with high accuracy, having mass errors below 5 ppm (Table S1) in the negative ion mode with respective retention times of ≈ 9 min for GMO and ≈ 10 min for OA in the extracted ion chromatogram (Figure 3B). Similarly, we also confirmed the membrane heterogeneity of HPs generated in the OA + MA reaction (Figure S6, Table S1). These results showed that the HP membranes generated in our experiment were indeed compositionally heterogeneous in nature. We also tried testing the membrane heterogeneity of the OA + GMO + OOH HP system. However, there was not sufficient HP pellet formation after centrifugation, likely because of the lower yield of HPs in this system compared to the other two heterogeneous systems (OA + GMO and OA + MA).

**Figure 3.**
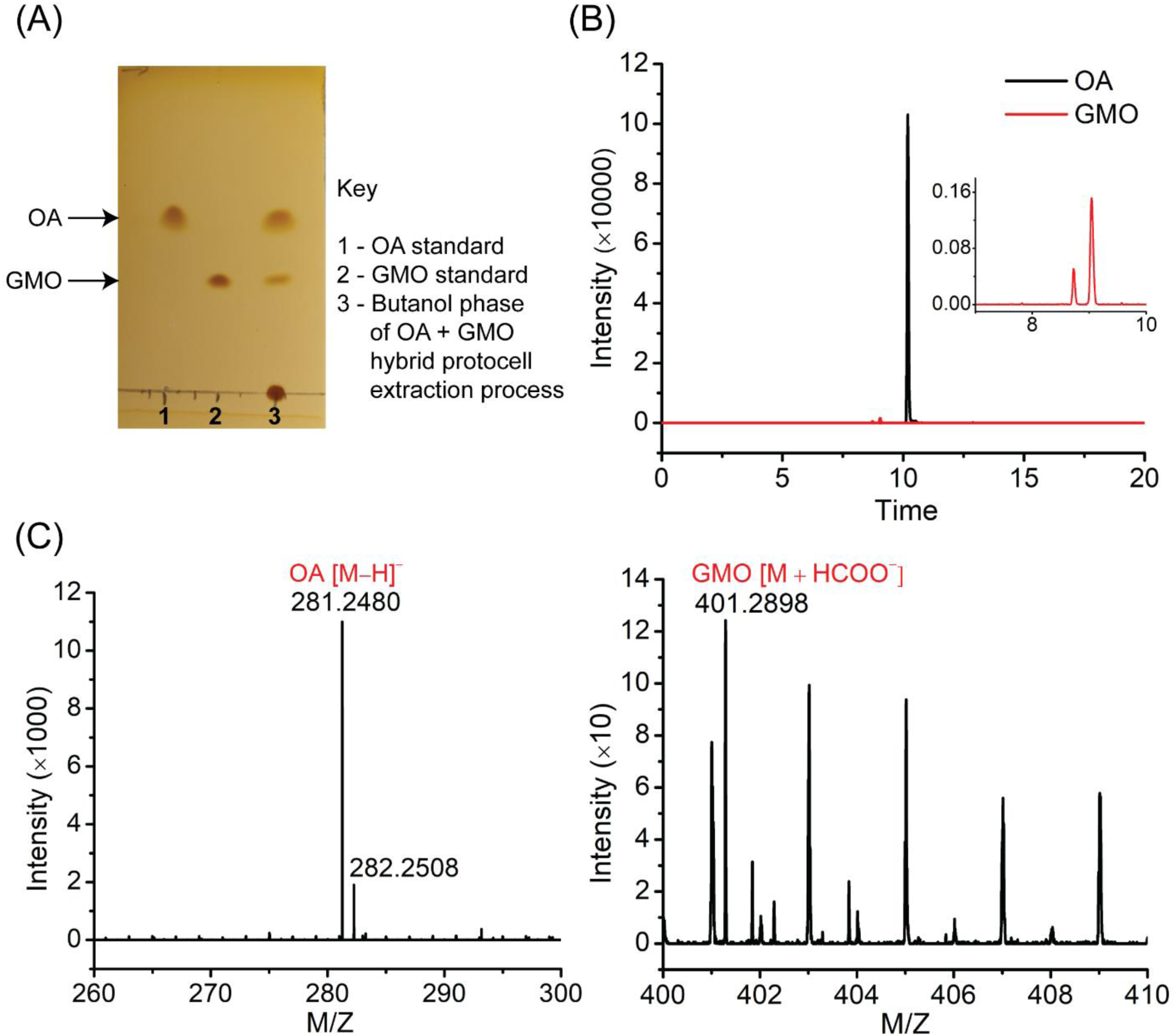
Confirming the heterogeneous nature of hybrid protocell membranes. After isolating the amphiphiles specifically from the OA + GMO (9 mM; 2:1 ratio) HP membrane, **(A)** the amphiphiles were detected with TLC, where lanes 1 and 2 represent OA and GMO standards, respectively, whereas lane 3 represents the amphiphiles present in the final butanol phase of the extraction process. TLC was performed with a silica stationary phase and toluene, chloroform, methanol (5:4:1 ratio) as a mobile phase with iodine staining. The presence of amphiphiles was further confirmed by mass spectrometry in negative ion mode. **(B)** The extracted ion chromatogram shows the peaks corresponding to GMO and OA ions with retention times of ≈ 9 min (see inset) and ≈ 10 min, respectively. **(C)** The corresponding mass spectra show the expected masses for GMO (401.2898), OA (281.2480), and the ^13^C isotope of OA (282.2508).

### Membrane heterogeneity affects the physicochemical properties of hybrid protocells

After establishing the formation of HPs with compositionally heterogeneous membranes, we investigated how increased membrane heterogeneity influences their properties, including HP formation efficiency (yield), size distribution, and the propensity to form multiple sub-compartments within a single HP structure. We also examined the impact of compositional heterogeneity on membrane dynamics, such as membrane order, lateral lipid diffusion, and membrane permeability. For these and all subsequent comparative analyses, OA-containing HPs were used as representatives of compositionally homogeneous HPs, whereas hybrid protocells containing both OA and GMO were used as representatives of compositionally heterogeneous HPs.

Overall, the formation efficiency of OA + GMO HPs was substantially higher than that of OA-only HPs. Quantitative analysis across four independent replicates for each type revealed approximately 1,867 OA + GMO HP structures, which is nearly threefold higher than the number of OA HPs observed (553). The size distributions of the two systems were also markedly different (Figures 4A and S7). OA + GMO HPs showed a greater tendency to form smaller structures, with nearly 70% of the population having areas below 40 µm², compared to only 32% of OA HPs falling within this size range (Figure S7). Notably, both OA and OA + GMO HPs exhibited multiple sub-compartments within individual HP structures, as demonstrated in Figure 4B. However, this feature was more pronounced in OA + GMO HPs, where approximately 28.5% of the total population contained two or more sub-compartments, compared to 20.2% in OA HPs (Figures 4C and S7).

**Figure 4.**
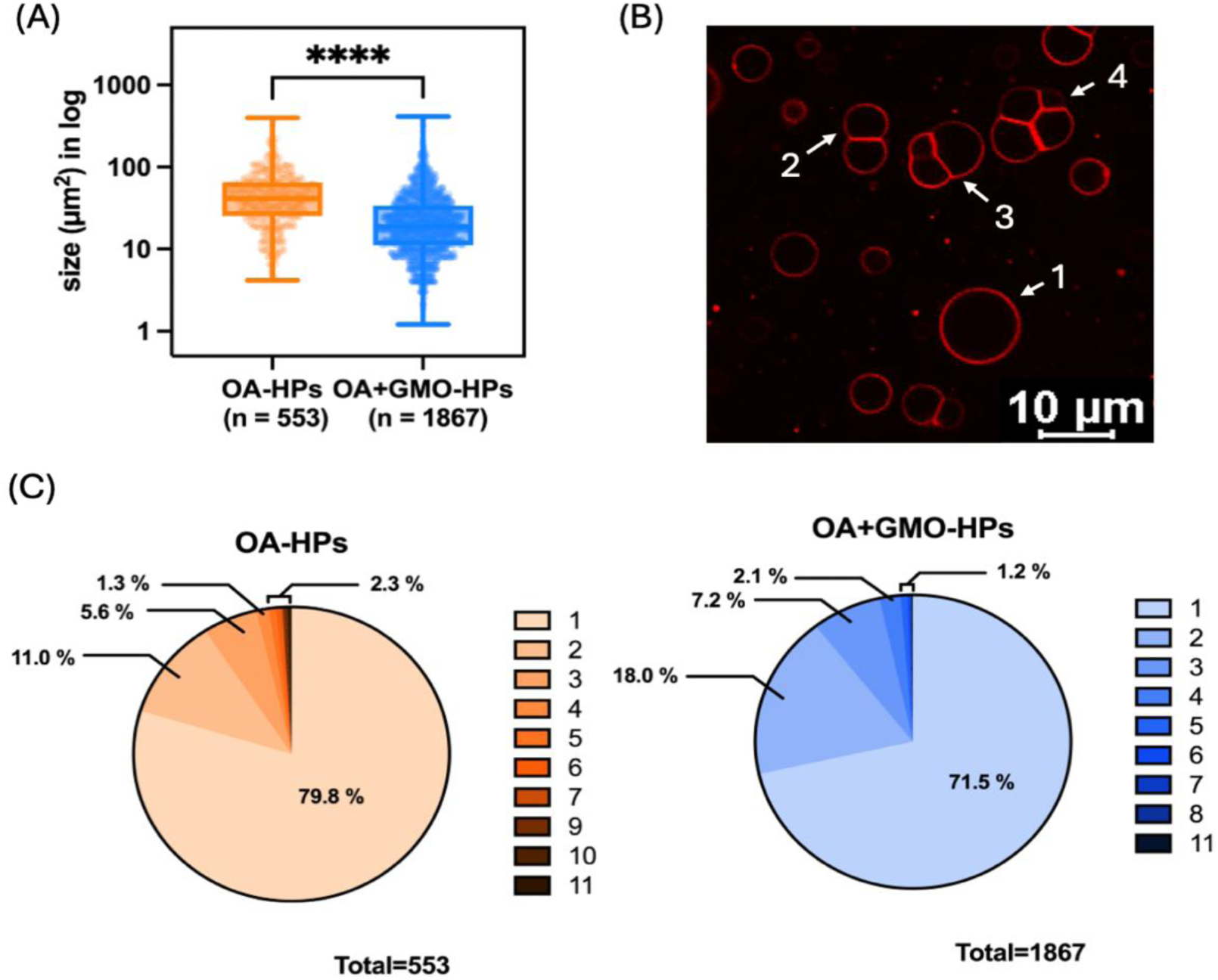
Comparison of size distribution and sub-compartment formation in homogeneous and heterogeneous hybrid protocells. **(A)** Box-and-whisker plots showing the size distribution of OA and OA + GMO HPs, calculated from images as the cross-sectional area of HP structures. Data points are shown alongside the box with center lines showing the medians, box limits indicating the 25^th^ and 75^th^ percentiles, and whiskers extending to the most extreme data points. Error bars represent standard deviation. The statistical significance is calculated using a two-tailed unpaired t-test, and **** represents p < 0.0001. **(B)** Image of OA + GMO HPs showing examples of hybrid protocell structures characterized as having 1, 2, 3, and 4 compartments. **(C)** Pie charts depicting the distribution of sub-compartments within individual HP structures for OA and OA + GMO systems. Each color represents the number of sub-compartments per HP in each system, as indicated in the legend. The total number of HPs with five or more sub-compartments is aggregated and presented as a single value.

The effect of heterogeneity on membrane order was studied using the 6-dodecanoyl-2-dimethylaminonaphthalene (Laurdan) dye, whose fluorescence properties depend on the polarity of its microenvironment, which, in turn, depends on membrane order when Laurdan is embedded in the membrane. A fluid membrane is more accessible to water molecules, which causes a red shift in the Laurdan emission spectrum (E_max_ = 490 nm) due to solvent relaxation, whereas a tightly packed membrane with limited access to water molecules results in a blue-shifted Laurdan emission spectrum (E_max_ = 440 nm). The fluorescence intensities at 440 nm and 490 nm are used to calculate Generalized Polarization (GP), which provides an estimate of membrane order, where a higher GP value indicates higher membrane order and vice versa. The OA (9 mM) and OA + GMO (9 mM; 2:1 ratio) HPs showed distinct Laurdan emission spectra with a significant difference around 440 nm (Figure 5A). The GP values of both OA and OA + GMO HP membranes were negative (Figure 5B), indicating a high degree of membrane disorder, increased water penetration, and loose lipid packing, which are characteristic features of membranes made of SCAs.^36^ However, the slightly higher GP value of OA + GMO HPs compared to that of OA HPs implies increased membrane order in OA + GMO HPs (Figure 5B). OA + GMO vesicles were also found to be more ordered than OA vesicles (Figure S8), consistent with earlier reports.^37^ The presence of GMO can increase the overall membrane order by filling void spaces in the membrane with its wedge-like shape, forming hydrogen bonds with carboxylate head groups of fatty acids, and creating small, ordered domains within the membrane. Interestingly, the membrane order was higher in OA vesicles compared to OA HPs, whereas no significant difference was observed between the membrane order of OA + GMO vesicles and HPs (Figure S8). This observation suggests that interactions between OA headgroups and amine moieties in the coacervate interfere with membrane order, and that incorporation of GMO, which, in addition to its other membrane-supportive roles, also reduces the density of these interactions, provides a remedy. We checked the localization of the Laurdan dye in the HP structure to verify that it was reporting on the membrane. Although PAH/ADP coacervates can independently encapsulate the externally added Laurdan, when they are surrounded by a membrane (OA or OA + GMO), the Laurdan dye preferentially goes to the membrane instead of the coacervate (Figure S9).

**Figure 5:**
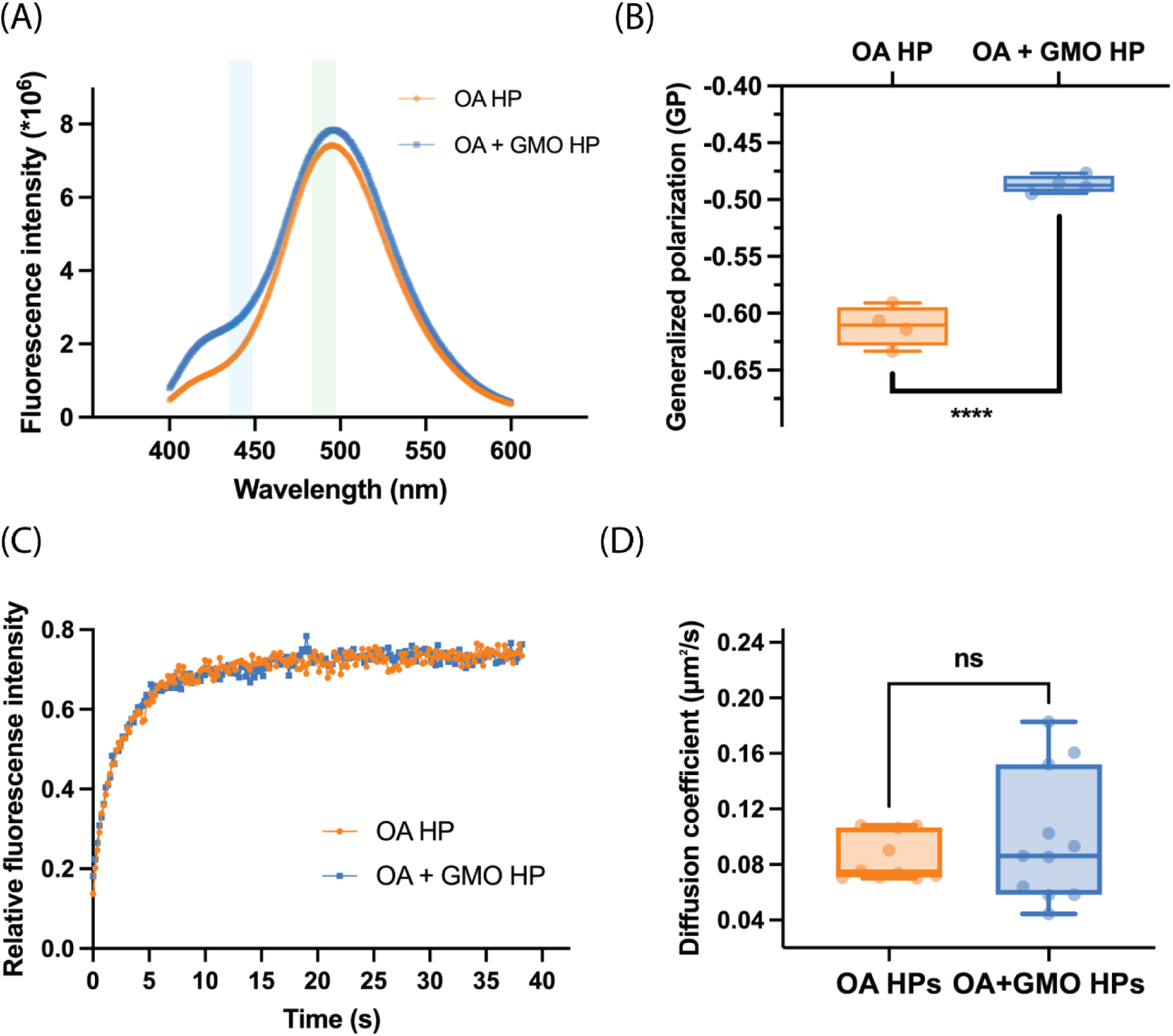
Effect of heterogeneity on the membrane order and the lateral diffusion of amphiphiles in the hybrid protocell membrane. **(A)** Laurdan emission spectra of OA (9 mM) and OA + GMO (9 mM; 2:1 ratio) HPs. **(B)** Generalized polarization (GP) values of OA and OA + GMO HPs based on Laurdan spectra. **(C)** Representative FRAP curves of R18 dye in the membranes of OA (9 mM) and OA + GMO (9 mM; 2:1 ratio) HPs. **(D)** Apparent diffusion coefficients of R18 dye as calculated from fluorescence recovery curves. In both (B) and (D), data is represented as a box and whisker plot, where data points are shown alongside the box with center lines showing the medians, box limits indicating the 25^th^ and 75^th^ percentiles, and whiskers extending to the most extreme data points. Error bars represent standard deviation. The statistical significance is calculated using a two-tailed unpaired t-test. For GP values, n = 4, and **** represents p < 0.0001. For apparent diffusion coefficient values, n = 9, and n.s. means not significant.

We also compared the rate of lateral diffusion of amphiphiles in the homogeneous and heterogeneous HPs by monitoring the fluorescence recovery after photobleaching (FRAP) of a single-chain lipophilic R18 dye. Despite the difference between the membrane order of homogeneous and heterogeneous HPs, their lateral diffusion coefficients were similar. Both OA and OA + GMO HP systems showed comparable fluorescence recovery curves (Figure 5C) with average diffusion coefficients of 0.08 µm^2^s^-1^ and 0.09 µm^2^s^-1^, respectively, where the difference between the two coefficients was statistically non-significant (Figure 5D). Although membrane order and the rate of lateral diffusion are generally inversely correlated, this relationship can be influenced by a variety of factors, like membrane composition, size, and shape of the membrane-forming molecules, and their interaction with each other. Moreover, these methods may focus on different regions of the membrane and operate over varying spatial and temporal scales, which can lead to non-consistent results.^47^

Subsequently, we compared the permeability of homogeneous and heterogeneous membranes to the externally added molecule. Four different solutes of varying sizes and charges were used in this study: fluorescein (332 g/mol, anionic, -2 charge), calcein (622 g/mol, anionic, -4 charge), FITC-Dextran 4k (∼4k Da, neutral), and Cy5-oligo RNA (U15, 5064 g/mol, anionic, - 15 charge). Each of these solutes readily accumulates within the coacervates when there is no membrane barrier, with fluorescence intensity ratios inside versus outside the coacervates (in/out) well above 1.0, indicating their preferential localization within the coacervates.^27^ The membrane permeability to these solutes was monitored for 30 minutes, 2 hours, and 24 hours. Despite the membrane disorder observed in the experiments of the previous section, both OA and OA + GMO HPs successfully restricted the penetration of large-sized and more charged solutes like calcein, FITC-Dextran 4k, and U15 RNA (Figure 6A), which was also reflected in their fluorescence intensity (in/out) ratios that remained below 0.5 (Figure 6B). However, some permeability to the small anionic molecule fluorescein was observed, with in/out intensity ratios approaching 1.0 within 24 hours. The impermeability of both types of membranes to externally added large neutral and anionic molecules is consistent with a prior report for HPs having PAH/ADP coacervate cores and all-OA membranes,^27^ and likely the consequence of the combined effects of molecular size and electrostatic repulsion from the negatively charged membrane, hindering solute diffusion across the membrane. These data show that the incorporation of the GMO in the HP membranes did not diminish their impermeability to these solutes. In the case of fluorescein, an interesting phenomenon was observed in the permeability behavior of these two systems. OA membranes displayed more uniform permeability across different structures, whereas OA + GMO membranes exhibited a differential permeability behavior within the population, with some HPs being permeable to fluorescein molecules while others restricting their entry (top panel in Figure 6A). This variation was particularly apparent at shorter time points (30 minutes and 2 hours) as fluorescein gradually penetrated the membranes. By 24 hours after fluorescein addition, penetration was largely complete for both OA and OA+GMO HPs (Figure 6B). The differential permeability observed in OA + GMO HPs is likely a consequence of subtle differences in membrane composition (i.e., OA:GMO ratio) and/or membrane organization (e.g., lamellarity, domain formation, etc.) between individual HPs within the population.

**Figure 6:**
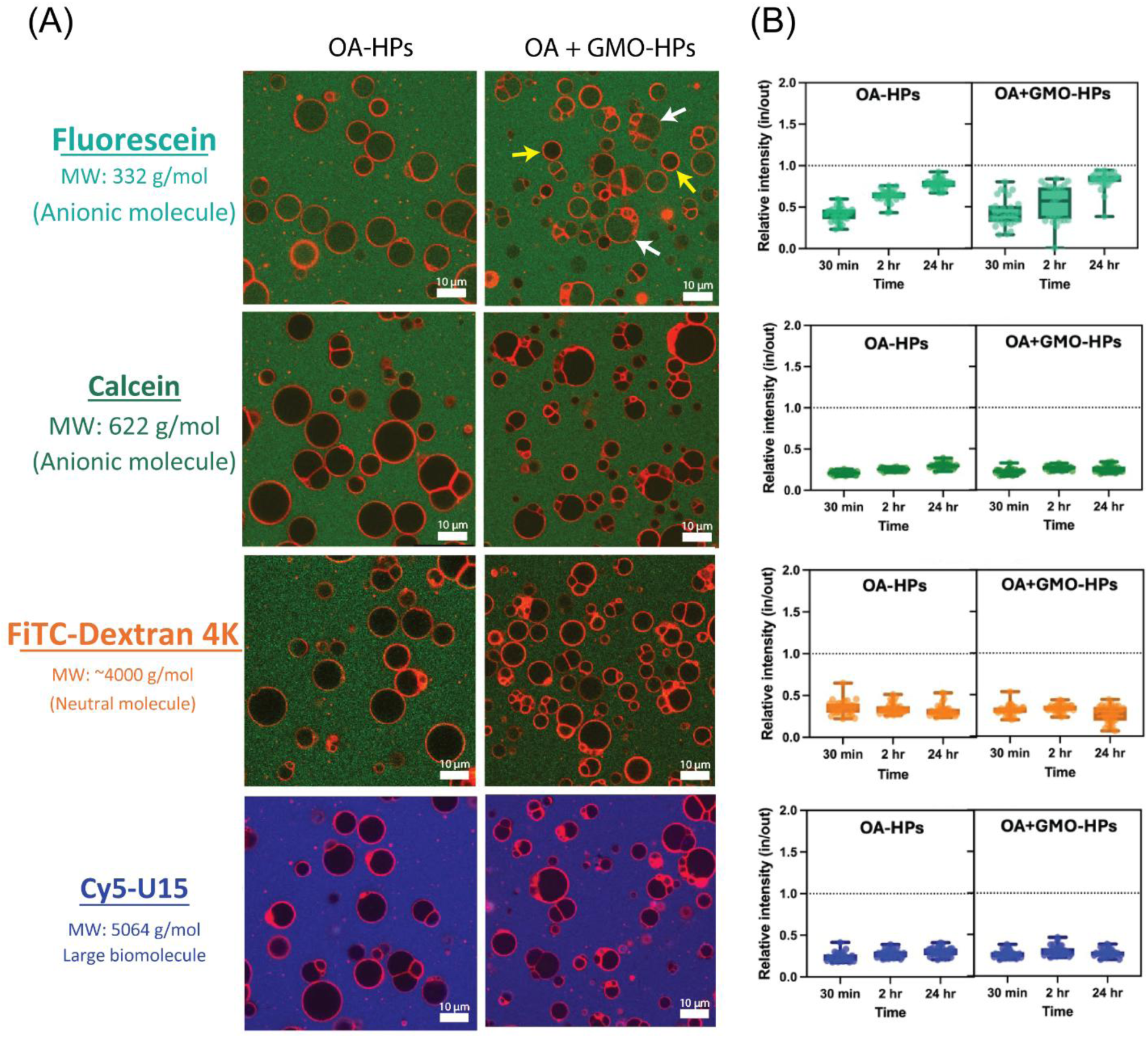
Effect of heterogeneity on the membrane permeability of hybrid protocells. **(A)** Confocal fluorescence images of OA-based hybrid protocells (OA-HPs) and OA + GMO-containing hybrid protocells (OA+GMO-HPs) incubated with various solutes for 2 hours. Solutes include Fluorescein (green, MW: 332 g/mol; anionic), Calcein (green, MW: 622 g/mol; anionic), FITC–Dextran 4K (green, MW: ∼4000 g/mol; neutral), and Cy5-U15 (blue, MW: 5064 g/mol; RNA oligomer, large biomolecule). Fluorescein permeable and impermeable hybrid protocells are shown by white and yellow arrows, respectively. Membranes are counterstained in red with R18 dye for all samples. **(B)** Quantification of solute permeability in OA-HPs and OA+GMO-HPs over time (30 minutes, 2 hours, and 24 hours), shown as inside versus outside (in/out) relative fluorescence intensity ratio for each solute.

### Effect of heterogeneity on pH stability of hybrid protocells

Prebiotic settings, such as hydrothermal vents and terrestrial hot springs, that are considered plausible sites for the origin of life, would have exhibited a wide range of pH conditions from highly acidic to strongly alkaline.^48,49^ To survive in such dynamic settings, early protocells would have needed to maintain structural integrity across varying pH levels. Fatty acid vesicles, which are commonly studied as model protocell systems, form only within a narrow pH range around their apparent pKa and are highly sensitive to pH changes.^50^ However, the incorporation of prebiotically plausible SCAs with polar, non-ionizable head groups, such as monoglycerides or fatty alcohols, has been shown to enhance the stability of fatty acid membranes mainly in the alkaline range,^31,33^ through hydrogen bonding with the carboxylate groups of fatty acids. We wanted to check whether such membrane heterogeneity also affects the pH stability of more complex assemblies, like HPs. Therefore, we compared the stability of homogeneous and heterogeneous HPs under more acidic and alkaline conditions relative to the initial pH at which they were formed. The OA (9 mM) and OA + GMO (9 mM; 2:1 ratio) HPs were prepared at pH 8.5 (± 0.1). Then the pH was systematically varied in the acidic or alkaline range by adding the specific volumes of HCl or NaOH solutions, respectively (see methods). We also tested the stability of OA (9 mM) and OA + GMO (9 mM; 2:1 ratio) vesicles (lacking a coacervate core) under the same conditions to delineate the effect of the presence of a coacervate on the stability of HPs.

During alkaline pH variations, OA HPs were surprisingly found to be stable at pH 10.6 (Figure 7A), which is significantly beyond the typical stability range for pure OA membranes.^31^ This increased stability is likely due to the enhanced ability of OA molecules to interact with the positively charged coacervate surface, because most of the OA molecules would be deprotonated and negatively charged at pH 10.6, which is above the apparent pKa of OA.^50^ However, pure OA vesicles that do not contain such a stabilizing coacervate core were disrupted at pH 10.6 (Figure S10). Furthermore, there was a formation of crystal-like aggregates, likely due to interactions between deprotonated OA molecules and excess Na^+^ ions added as NaOH during pH adjustment. This crystallization was markedly reduced, and some vesicles were also observed when OA vesicles were prepared in the dilute phase of PAH/ADP coacervates (Figure S11), highlighting the stabilizing effect provided by free PAH and ADP molecules in the surrounding medium. OA + GMO vesicles were relatively stable at pH 10.6 (Figure S10), which is consistent with the stabilizing role of GMO as reported earlier.^31^ Together, these results suggested that the enhanced alkaline stability of HPs was likely due to the combined effect of various stabilizing factors, such as the coacervate core, coacervate-forming molecules in the surrounding medium, and the presence of GMO in the case of heterogeneous membranes.

**Figure 7.**
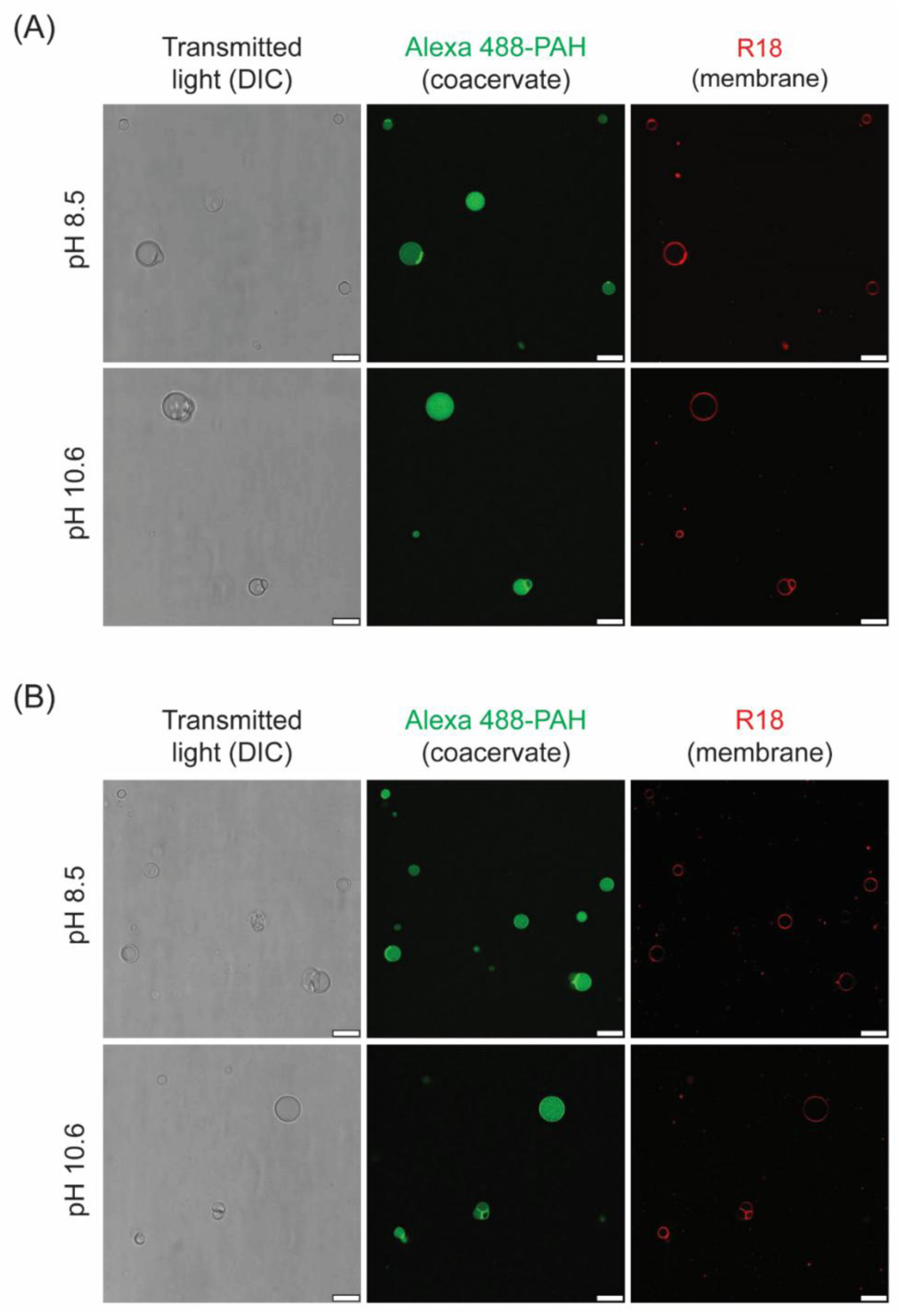
Stability of hybrid protocells at varying alkaline pH. **(A)** OA (9 mM) and **(B)** OA + GMO (9 mM; 2:1 ratio) HPs were prepared at pH 8.5, and the pH was increased to 10.6 by adding 1 M NaOH solution. Differential Interference Contrast (DIC, left panels) images show the actual morphology of hybrid protocells without staining, whereas the membrane and coacervate parts of the hybrid protocell are differentially labeled with Octadecyl Rhodamine B (R18) (red color, right panels) and Alexa Fluor 488-PAH (green color, middle panels), respectively. Fluorescence images have been pseudo-colored and contrast-adjusted for better visualization. All scale bars are 10 µm.

Subsequently, we investigated the stability of HPs under acidic conditions by systematically lowering the pH from 8.5 to 7.4 and 6 through controlled HCl addition (see Methods). Although both OA and OA + GMO HPs exhibited reduced stability upon acidification relative to their initial pH, in terms of HP abundance and morphology, OA + GMO HPs showed slightly improved stability compared to OA HPs at pH 7.4 (Figure 8A, middle panels). However, at pH 6, neither system was able to maintain structural integrity, resulting predominantly in aggregated structures rather than well-defined HPs. This destabilization was even more pronounced for the corresponding vesicles, which formed aggregates at both pH 7.4 and 6 (Figure S12). These observations indicate a stabilizing effect of the coacervate core on HP membranes under acidic conditions. Also, our results are consistent with the known pH-dependent self-assembly behavior of fatty acids, where they transition from membrane bilayers to hydrophobic oil droplets at pH values below their apparent pKa.^51^ It also explains the observed aggregatory nature of fatty acid-based membranes of HPs and vesicles in our experiments.

**Figure 8.**
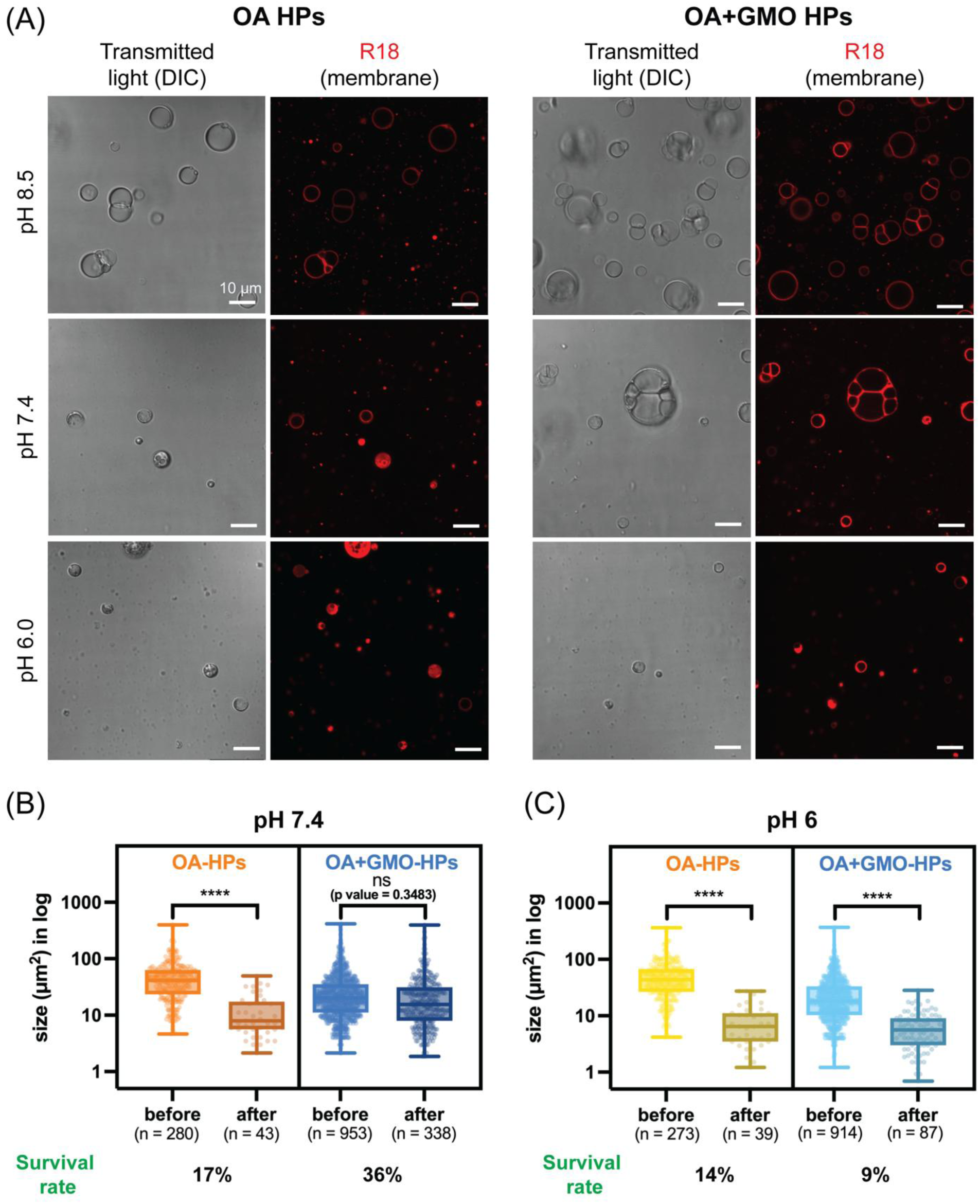
Stability of hybrid protocells under acidic pH fluctuations. OA (9 mM) and OA + GMO (9 mM; 2:1 ratio) HPs were prepared at pH 8.5, and the pH was decreased to 7.4 and 6.0 by adding 1 M HCl solution. **(A)** Transmitted light (DIC) and fluorescence microscopy images showing the structural changes in HPs upon pH change. HP membranes are labeled with R18 dye for fluorescence imaging. The presence of a coacervate was confirmed as a dense structure compared to the surrounding medium. Statistical analysis of the change in HP size at **(B)** pH 7.4 and **(C)** pH 6. In both (B) and (C), the data are represented as box and whisker plots, with details regarding the graphs and statistical analyses the same as those in the earlier figure legends.

Given the comparatively improved stability of OA + GMO HPs at pH 7.4, we decided to perform a detailed quantitative investigation of the impact of these pH fluctuations on HP yield, size retention, and sub-compartment heterogeneity (the presence of multiple sub-compartments within a single HP structure). Consistent with the microscopy observations, at pH 7.4, OA + GMO HPs retained 328 intact structures out of 920 initially observed at pH 8.5, corresponding to a survival rate of 36%. In contrast, OA HPs showed a more pronounced loss, with HP counts decreasing from 280 to 43 (15% survival). Further acidification to pH 6 resulted in a substantial decline in HP yield for both systems. OA HPs decreased from 273 to 39 structures (14% survival), while OA + GMO HPs decreased from 852 to 80 structures (9% survival).

The stability pattern observed in HP yield was also reflected in their size and sub-compartment heterogeneity. OA HPs exhibited a marked reduction in both mean size and size distribution when the pH was lowered from 8.5 to 7.4. In contrast, OA + GMO HPs largely preserved these parameters under the same conditions (Figures 8B and S13). At pH 6, both systems showed significant reductions in overall size and size distribution (Figures 8C and S13). Notably, OA + GMO HPs maintained sub-compartment heterogeneity more efficiently than OA HPs at both pH 7.4 and 6 (Figure S14).

Overall, HPs having mixed-SCA membranes displayed enhanced resistance to acidic pH fluctuations, not only in terms of survival but also in retaining key structural features such as size and sub-compartment heterogeneity. This improved performance can be attributed to the combined effects of membrane compositional heterogeneity and the stabilizing influence of the coacervate core.

## CONCLUSIONS

This study demonstrates that hybrid protocells with compositionally heterogeneous membranes can be readily generated using mixtures of SCAs with diverse head groups and alkyl chain lengths, which are commonly used to prepare model protocell membranes. By incorporating SCA mixtures that provide increased chemical complexity in SCA membrane-coated coacervate systems, these hybrid protocell models become more realistic and versatile for studies pertaining to the origin of life and synthetic biology. We found that membrane compositional heterogeneity strongly influences the self-assembly behavior and physicochemical properties of HPs. Overall, heterogeneous HPs exhibited higher yields and a greater propensity for multi-compartment formation compared to homogeneous HPs. It reflects the robustness of heterogeneous HPs and their potential to generate sub-compartments with well-defined membrane boundaries. Such compartmentalization could enable the spatial segregation of distinct protometabolic reactions, thereby recapitulating fundamental principles of compartment-specific reaction networks observed in contemporary cells.

Compositional heterogeneity also influenced membrane dynamics, where HPs with heterogeneous membranes exhibited enhanced order and showed population-level variability in the permeability to a small anionic molecule, which may have significant implications for protocell evolution. Such variability could enable competition among protocells regarding nutrient uptake and protection of genetic material and other essential metabolites from harmful external substances. Thus, some protocells may gain a selective advantage over others under environmental pressures such as resource scarcity or exposure to toxic compounds. In contrast, such selective pressures would likely be ineffective in homogeneous systems, where all protocells exhibit similar behavior.

Our pH stability studies further reveal that in complex systems, such as HPs, overall stability is governed by multiple factors, including the coacervate core, free coacervate-forming molecules in the surrounding medium, and membrane heterogeneity. While both homogeneous and heterogeneous HPs were stable under alkaline conditions, the latter system showed enhanced stability under acidic pH fluctuations, as reflected in their overall yield and retention of size and sub-compartment heterogeneity. This suggests that heterogeneous protocells would be more resilient to pH fluctuations on early Earth than their homogeneous counterparts.

The work presented here further supports the growing realization that increased chemical complexity in prebiotic model systems can be beneficial. SCA membranes can be stabilized by mixed-lipid composition,^30,39^ by interactions with small molecule solutes such as nucleotides and peptides,^52–54^ and also by interacting with an internal coacervate droplet that serves as a model cytoplasm.^27^ The HPs described here combine these nonadditive effects in one system. Future studies could explore more prebiotically plausible molecular combinations to create HPs, including amphiphiles with shorter chains (C ≤ 10) and/or additional headgroup chemistries for the membranes,^35,55–59^ as well as simple oligopeptide and/or oligonucleotide mixtures^12,60–63^ for their coacervate components. Additionally, it would be interesting to investigate how membrane heterogeneity governs the selective uptake of molecules from a mixture in the surrounding medium, which could influence key protocell functions such as protometabolism,^61,64,65^ ribozyme-mediated catalysis,^66,67^ or the replication of genetic material.^68,69^

## MATERIALS AND METHODS

### Materials

Oleic acid (OA), Myristoleic acid (MA), Monoolein (GMO), and Oleyl alcohol (OOH) were purchased from Nu-Chek Prep. Poly(allylamine hydrochloride) (PAH, MW 17.5 kDa), Adenosine 5’-diphosphate (ADP) disodium salt, bicine, tris (Trizma Base), fluorescein, calcein, Cy5-labeled U15 (MW 5 kDa), and FITC-Dextran (MW 4 kDa) were purchased from Sigma-Aldrich. 6-Dodecanoyl-2-Dimethylaminonaphthalene (Laurdan), Octadecyl Rhodamine B chloride (R18), and Alexa Fluor 488 NHS ester were purchased from Invitrogen (Thermo Fisher Scientific). The Alexa Fluor 488 was used for labeling PAH molecules as follows: A total of 10 µl of Alexa Fluor 488 dye solution (10 mg/ml in DMSO) was added to 1 ml of 1 mg/ml PAH in 10 mM HEPES buffer pH 7.6, with 2 µl addition at a time with pipette mixing. This PAH and dye mixture was further moderately mixed for 1 hour at RT using a vortex shaker, allowing the labeling reaction to occur. The free dye was separated from the PAH-labeled dye using an Amicon Ultra Centrifugal Filter 10 kDa MWCO (MilliporeSigma).

### Preparation of hybrid protocells

The hybrid protocells were prepared by a modified gentle hydration method. First, amphiphile stock solutions were prepared in methanol (250 mg/ml for OA, 200 mg/ml for MA, and 100 mg/ml each for GMO and OOH, respectively). Appropriate volumes of the amphiphile stock solutions were added to the glass tube containing chloroform so that the total amphiphile concentration in the final reaction (after hydration) would be 9 mM. Lipid films for homogeneous hybrid protocells were prepared using 9 mM OA, whereas those for heterogeneous hybrid protocells were prepared using the following amphiphile combinations: OA + GMO (9 mM; 2:1 ratio), OA + GMO + OOH (9 mM; 4:1:1 ratio), and OA + MA (15 mM; 1:9 ratio). To stain the hybrid protocell membrane, the appropriate volume of R18 dye stock solution (1 mg/ml in methanol) was added during lipid film formation to get a 9 µM final dye concentration (1:1000 molar ratio of dye to total lipid). This amphiphile mixture was dried under a vacuum for 6 hours to completely evaporate the organic solvent and prepare a lipid film.

PAH and ADP stock solutions were prepared in nanopure water (resistivity ≍ 18.2 MΩ·cm at 25°C) with a concentration of 5 wt.% and 100 mM, respectively, and their pH was adjusted to 7.4 using 1 M NaOH. The PAH stock solution had a net charge of + 563 mM, while the ADP stock solution had a net charge of - 300 mM (considering - 3 charges per ADP molecule at pH 8.5). 1 M bicine buffer was also prepared in nanopure water, and its pH was adjusted to 8.5 using 1 M NaOH. The PAH/ADP coacervates were prepared by adding solvent and stock solutions in the following order to get the desired final concentration of each component: nanopure water, bicine buffer (100 mM, pH 8.5), ADP (6.67 mM; - 20 mM charge concentration), and PAH (+ 20 mM charge concentration). The PAH was premixed with Alexa Fluor 488-labeled PAH to stain the coacervates (0.5 or 1 µl of dye solution (depending on the batch) for 500 µl of coacervate solution). After the addition of the final component (PAH), the solution immediately became turbid, indicating the coacervate formation. The solution was gently pipette mixed 3 times and immediately added to the lipid film.

After hydrating the lipid film with the PAH/ADP coacervate solution, the mixture was incubated at 45°C for 30 minutes with gentle vortex mixing before incubation, after 15 minutes, and at the end of the incubation. Hybrid protocell formation was confirmed by confocal imaging (Leica TCS SP5) using a 63×/1.40 NA oil immersion objective. The membrane and the coacervate components were differentially visualized by exciting R18 dye at 543 nm and Alexa Fluor 488-labeled PAH at 488 nm, respectively. The overall morphology of hybrid protocell structures was checked using DIC microscopy. The same protocol was followed for generating hybrid protocells in all the experiments unless otherwise mentioned. OA + MA hybrid protocells were generated using 100 mM tris pH 8, considering the lower apparent pKa of MA.

### Confirming the membrane heterogeneity of hybrid protocells

OA + GMO (9 mM; 2:1 ratio) and OA + MA (15 mM; 1:9 ratio) hybrid protocells were prepared as described above, except without the addition of Alexa Fluor 488-labeled PAH to the coacervates. The solutions were scaled up to obtain a sufficient number of hybrid protocell structures. As control reactions, OA + GMO and OA + MA vesicles were prepared under the same conditions except that the lipid films were hydrated with respective buffers instead of a PAH/ADP coacervate solution. The hybrid protocells were pelleted by centrifugation at 15,700 RCF for 20 min. This pellet was absent in the control reactions containing only vesicles. After centrifugation, the supernatant (containing vesicles and free amphiphiles) was carefully removed, and the hybrid protocell pellet was resuspended in 100 mM bicine buffer. To this solution, an equal volume of butanol was added and vigorously mixed through vortexing. The solution was then centrifuged at 15,700 RCF for 1 min to separate the butanol layer (containing amphiphiles) and the aqueous layer (containing coacervate-forming molecules). The upper butanol layer was carefully transferred to a new microfuge tube, and the butanol extraction process was repeated. The butanol layers from both steps were pulled into a single tube, and the butanol was then completely evaporated. The resultant lipid film was redissolved into a smaller volume of butanol to achieve a detectable concentration of amphiphiles. This butanol solution was first analyzed by thin-layer chromatography (TLC) using a normal-phase silica plate as a stationary phase and the mixture of toluene, chloroform, and methanol in a 5:4:1 ratio (v/v) as a mobile phase. The amphiphile spots were detected by iodine staining. The presence of amphiphiles was further confirmed by liquid chromatography-mass spectrometry (LC-MS) using a Nexera 40 HPLC system (Shimadzu) coupled to a ZenoTOF 7600 mass spectrometer (Sciex) by electrospray ionization in negative mode using an IonDrive Turbo V ion source. Chromatographic separation was performed by injecting 2 µL of sample (in methanol) into a reversed-phase C18 column (Waters, CORTECS 2.1 × 100 mm, 1.6 µm particle size) and eluting at 55°C with a 0.25 ml/min flow rate using a mobile phase containing Solvent A (60% acetonitrile in water (v/v) + 0.1% formic acid (v/v)) and Solvent B (100% acetonitrile + 0.1% formic acid (v/v)) in a gradient mode over 20 minutes as follows: 0-2 min, 2% B; 5 min, 75% B; 11 min, 85% B; 12 min, 98% B; 17-20 min; 2% B. MS1 and MS2 data were acquired using flowing parameters: declustering potential (DP) = - 60 V, ion spray voltage (IS) = - 4500 V, curtain gas (CUR) = 35 psi, nebulizer gas (GS1) = 50 psi, heater gas 2 (GS2) = 50 psi, and heater temperature (TEM) = 500 °C. The mass error was calculated using the following formula.

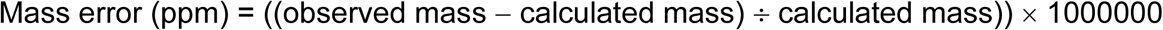

### Quantification of the size and multi-compartment formation efficiency of hybrid protocells

OA (9 mM) and OA + GMO (9 mM; 2:1 ratio) hybrid protocells were prepared according to the protocol described above, except that Alexa Fluor 488-labeled PAH was omitted from the coacervate phase. Microscopic imaging was performed using a Leica TCS SP5 confocal microscope equipped with an HCX PL APO CS 63.0×1.40 OIL UV objective. To visualize the hybrid protocell membranes, R18 dye was incorporated at a concentration of 5.468 µM (1:1646 dye-to-total lipid ratio) and excited using a 543 nm laser.

For each sample, ten images were captured at random locations. Randomness was ensured by moving the stage to a new field of view without pre-screening for specific structures, followed by minor re-focusing before acquisition. Once collected, the images were processed and analyzed using Fiji (ImageJ) software to determine the size distribution and the number of internal compartments within individual hybrid protocell structures.

For each hybrid protocell structure, the cross-sectional area was determined by manually defining the structure boundaries using either "oval selections" or "polygon selections," depending on the specific morphology of the hybrid protocell, in order to ensure that the structure was captured accurately for size distribution analysis.

It is important to note that because the majority of these hybrid protocells exhibited complex internal multi-compartmentalization, calculating total volume based solely on the diameter or area we obtained from the focused plane is inherently limited. Furthermore, the acquired images did not account for additional compartments that may exist above or below the focal plane. Consequently, the data presented herein represent a 2D quantification of size and visible compartment heterogeneity within the primary plane of focus.

### Estimating the membrane order of hybrid protocells and vesicles

OA (9 mM) and OA + GMO (9 mM; 2:1 ratio) hybrid protocells and vesicles were prepared without adding R18 and Alexa Fluor 488-PAH dyes to avoid any potential interference with the Laurdan fluorescence. To these solutions, an appropriate volume of Laurdan dye stock solution (900 µM in methanol) was added to get a 9 µM dye concentration in the final reaction volume (1:1000 ratio of dye to lipid). Then, the solutions were incubated for 15 minutes to allow the incorporation of Laurdan into the membrane. Laurdan fluorescence was measured using a fluorimeter (Jobin Yvon model FL3-21) by exciting the sample at 370 nm and recording the emission spectrum from 400 to 600 nm. Both excitation and emission slit widths were set to 2 nm. Spectra for control reactions (without Laurdan dye) were recorded using identical parameters and used for background signal subtraction. The Generalized Polarization (GP) values were calculated using the following formula, where ‘I’ indicates intensity at a specific wavelength.

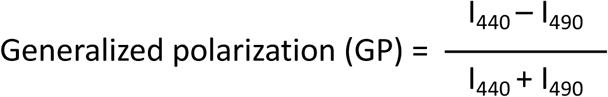

The GP values were calculated from four independent replicates and reported as mean ± standard deviation. The statistical significance was calculated using a two-tailed unpaired t-test.

The localization of Laurdan dye into a hybrid protocell membrane was confirmed by visualizing the samples under a fluorescence microscope with excitation at 405 nm and emission from 450 nm to 550 nm. The PAH/ADP coacervate sample (40 mM total charge; 1:1 charge ratio) containing Laurdan was also imaged using the same settings.

### Fluorescent recovery after photobleaching

FRAP studies were carried out using a FRAP module in a Leica TCS SP5 confocal microscope equipped with an HCX PL APO CS 63.0×1.40 OIL UV objective. FRAP experiments were conducted on OA (9 mM) and OA + GMO (9 mM; 2:1 ratio) hybrid protocell membranes containing R18 dye (5.468 µM; 1:1646 ratio of dye to total lipid) using excitation at 543 nm. A circular area of interest, measuring 2 µm, in the membrane was bleached using lasers (458, 476, 488, 514, 543, and 633 nm) at 100% power. FRAP recovery was monitored by acquiring 10 frames before bleaching, 15 frames during photobleaching, and 200 frames post-bleaching at every 0.19 s. Background noise was corrected by measuring fluorescence intensity in ROIs with all lasers turned off while keeping the respective photomultiplier tubes on. A reference ROI was also acquired to account for any photobleaching effects during normal imaging with the 543 nm laser. The apparent diffusion coefficients were calculated according to the protocol detailed in our previous work.^27^ The FRAP assay was conducted using 3 independent reaction replicates, with at least 3 data points obtained for each replicate.

### Membrane permeability of hybrid protocells

Permeability studies were performed using confocal microscopy (Leica TCS SP5) with an HCX PL APO CS ×63.0/1.40 NA oil UV objective. To conduct the permeability study, 20 µL of OA (9mM) and OA + GMO (9 mM; 2:1 ratio) hybrid protocell solution was added to separate coverslips, immediately followed by the addition of 1 µL of the desired solute to achieve final concentrations of 5 µM each for fluorescein, calcein, and FITC-Dex 4k, and 10 µM for Cy5-U15 RNA. At least five images (four corners and one in the middle of the sample drop) were acquired for hybrid protocell samples at the designated time points (30 minutes, 2 hours, and 24 hours) using confocal microscopy. The background signal was obtained using identical imaging settings from samples without solute addition. ImageJ (Fiji) was used to quantify fluorescence intensities inside and outside the hybrid protocell structures, with the outside intensity calculated as the average of five regions of interest (ROIs) positioned at the four corners and the center of each image. Average intensity ratios were calculated after subtracting background intensity from control samples (no fluorescent solute added). All experiments were performed in triplicate.

### pH stability of hybrid protocells and vesicles

For checking the stability of hybrid protocells in the alkaline range, the initial pH of 8.5 was increased to 10.6 using 1 M NaOH solution. The volume of NaOH to be added to make this pH change was first optimized with 100 mM bicine buffer pH 8.5, where the addition of 8.2 µl of 1 M NaOH to 200 µl of this buffer brought the pH to around 10.6, which was checked with a pH meter. The reproducibility of this NaOH-induced pH change was further confirmed with three independent replicates, which showed an average final pH of 10.58 ± 0.11 (s.d.) after NaOH addition. The same volume of NaOH (8.2 µl) was added to 200 µl of OA (9 mM), and OA + GMO (9 mM; 2:1 ratio) hybrid protocell solutions, followed by gentle pipette mixing, and the pH change was confirmed using pH indicator strips. The stability of hybrid protocell structures was checked with fluorescence and DIC microscopy, where the membranes were visualized by staining with R18 dye, as mentioned earlier. The same procedure was followed to check the alkaline stability of OA (9 mM) and OA + GMO (9 mM; 2:1 ratio) vesicles prepared in 100 mM bicine buffer and the dilute phase of PAH/ADP coacervates, which was obtained by centrifuging the coacervate solution at 15,700 RCF for 20 minutes.

For testing the acidic stability of OA and OA + GMO hybrid protocells, the initial pH of 8.5 was decreased to 7.4 using a 1 M HCl solution. The volume of HCl to be added to make this pH change was first optimized with 100 mM bicine buffer pH 8.5, where the addition of 2.25 and 2.6 µl of 1 M HCl to 50 µl of this buffer brought the pH to around 7.4 and 6, respectively, which was checked with a pH meter. The acid stability of hybrid protocells was then evaluated using a Leica TCS SP5 confocal microscope with both fluorescence and differential interference contrast (DIC) channels. For the stability assays, 50 µL of the HP solution was placed onto a coverslip equipped with a silicone spacer (Invitrogen, P18174) and sealed with a second coverslip to prevent evaporation. Initial "random" imaging was performed by translating the stage to new fields of view without pre-screening for specific structures, followed by minor re-focusing before acquisition. After the initial imaging, the top coverslip was carefully removed using fine-tipped tweezers. Any residual liquid on the top coverslip was recovered and transferred back to the sample. The optimized volume of 1 M HCl (2.25 µL or 2.6 µL) was added to the solution, followed by gentle pipette mixing six times. The samples were re-sealed and allowed to equilibrate for 10 minutes before a second set of ten random images was captured. After completing the image acquisition, final pH values were verified using pH indicator strips (pH ranges 6.5–10 and 0–14) to confirm that the targeted changes in the pH levels (pH 7.0–7.4 after 2.25 µL HCl addition, and pH ≤ 6.0 after 2.6 µL of HCl addition) were successfully achieved.

Digital image processing and analysis were performed using Fiji (ImageJ). To ensure objective and consistent quantification across all samples, HP structures were identified and counted based on three rigorous criteria:

1. Only structures entirely contained within the imaging frame were included in the analysis; partial structures overlapping the frame boundaries were excluded to ensure accurate area and size measurements.
2. To differentiate HPs from empty vesicles or lipid aggregates, each structure was cross-verified using the differential interference contrast (DIC) channel. Only objects exhibiting the characteristic high refractive index associated with a dense coacervate interior were recorded.
3. For samples analyzed after acidification, "surviving" hybrid protocells were defined by their membrane morphology. Structures were excluded from the final count if membrane aggregation or "patchiness" exceeded 50% of the total membrane surface area.

These criteria were applied to determine both the total survival counts and the distribution of internal compartments within individual HP structures. Size distributions and compartment heterogeneity were subsequently analyzed and visualized using Excel and GraphPad Prism 9.

## ACKNOWLEDGEMENTS

This project was supported by the NASA Exobiology Grant No. 80NSSC22K0553. Partial support was also provided by the Huck Institutes of the Life Sciences at Penn State University through a Huck Innovative and Transformational Seed Grant (HITS). M.P.J. was supported by an appointment to the NASA Postdoctoral Program at the Pennsylvania State University, administered by Oak Ridge Associated Universities under contract with NASA. Mass spectrometric analyses were performed at the Penn State Proteomics and Mass Spectrometry Core Facility, University Park, PA (RRID:SCR_024462), and we acknowledge Sergei Koshkin for collecting the data shown in Figures 3 and S6. The authors would also like to acknowledge the Huck Institutes’ Microscopy Core Facility (RRID:SCR_024457), and Ella Mullikin for her assistance with imaging.

