## Supplementary information for "Increasing the compositional heterogeneity of single-chain amphiphile membranes supported by coacervate cores alters stability and properties of the hybrid protocells"

Academy of Scientific and Innovative Research (AcSIR), Ghaziabad 201002, India

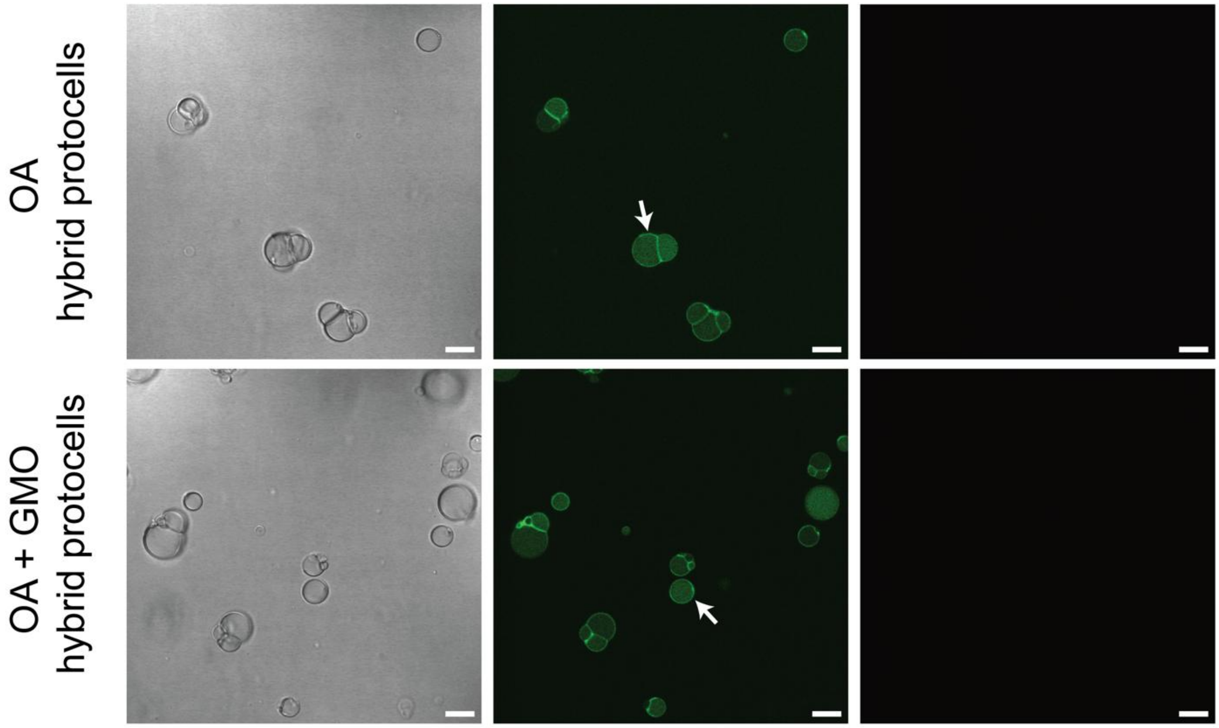


**Figure S1. Preferential localization of Alexa Fluor 488-labeled PAH on the membrane-coacervate interface is not due to its interaction with the membrane dye.** OA and OA + GMO hybrid protocells were generated without the membrane-staining R18 dye, and only the PAH/ADP coacervate part was stained with Alexa Fluor 488-labeled PAH (middle panels). White arrows indicate the preferential localization of Alexa Fluor 488-labeled PAH on the membrane-coacervate interface, even in the absence of R18 dye (right panels; no R18 fluorescence). Transmitted light, specifically differential interference contrast (DIC) images (left panels), show the actual morphology of hybrid protocell structures. The scale bar is 10 μm.

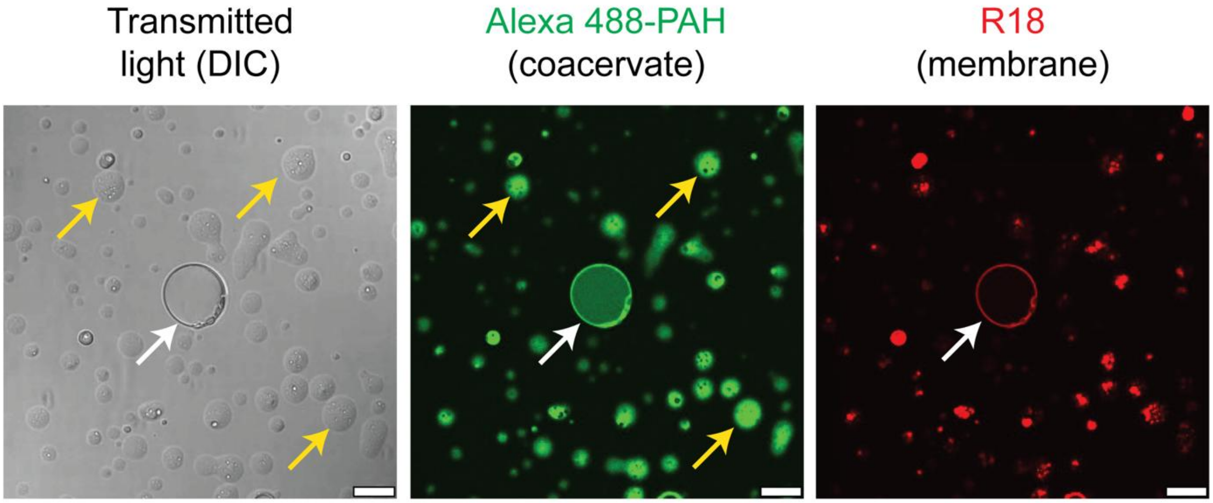


**Figure S2. Testing the hybrid protocell formation at a lower total amphiphile concentration.** 3 mM OA was used (instead of 9 mM OA) for generating hybrid protocells. The coacervate concentration was kept the same (40 mM total charge; 1:1 charge ratio), and they were prepared in 100 mM bicine, pH 8.5. The white arrow indicates hybrid protocells, whereas the yellow arrow indicates coacervates encapsulating amphiphiles. The membrane and the coacervate were stained with R18 and Alexa 488-PAH, respectively. The scale bar is 10 μm.

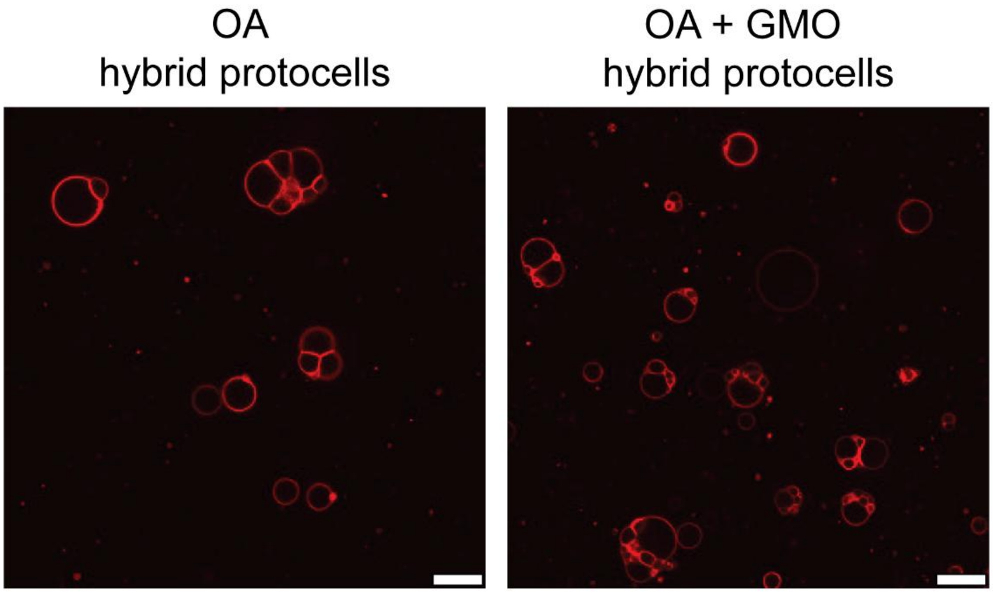


**Figure S3. OA and OA + GMO hybrid protocell membranes look similar under the microscope.** Both OA (9 mM) and OA + GMO (9 mM; 2:1 ratio) hybrid protocell membranes show similar morphology when observed under the confocal microscope. Both hybrid protocell membranes were stained with R18 dye. The scale bar is 10 μm.

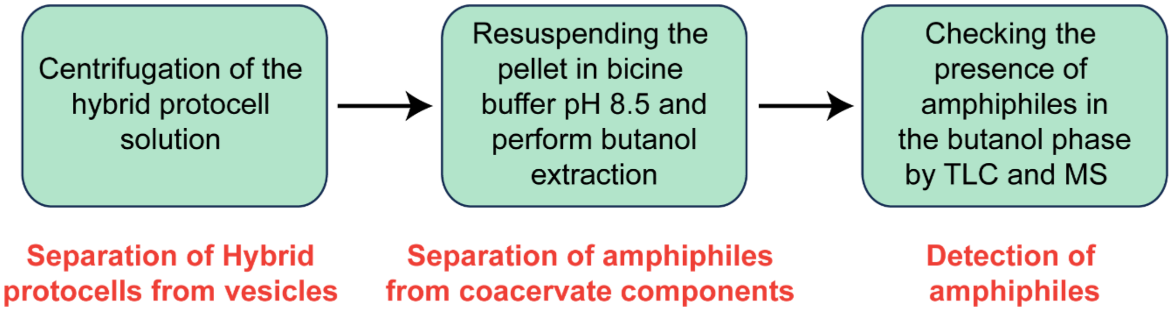


**Figure S4. Overview of the protocol developed for confirming the membrane heterogeneity of hybrid protocells.**

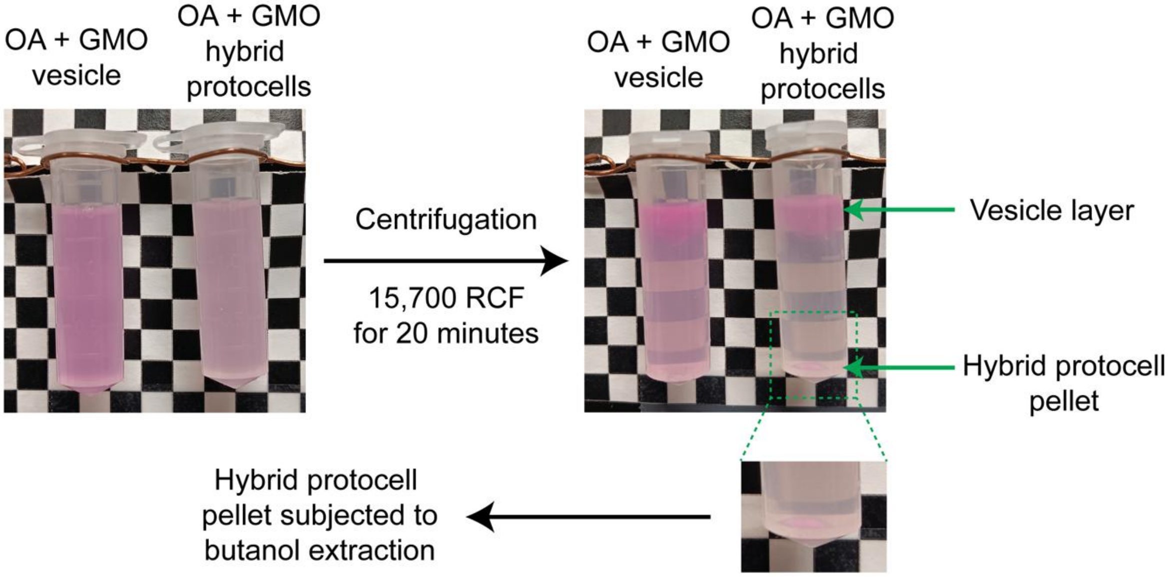


**Figure S5. Appearance of the hybrid protocell and vesicle sample before and after centrifugation.** Both hybrid protocells and vesicles were prepared using the same amphiphile concentration (OA + GMO; 9 mM, 2:1 ratio), but hybrid protocells contain a coacervate core, which is absent in vesicles. Both samples were added with amphiphilic R18 dye to track membrane structures. Initially, the vesicle sample appears as a typical uniform translucent solution, whereas the hybrid protocell sample appears as a whitish turbid solution. After centrifugation, vesicles float on the surface because of their buoyancy, whereas hybrid protocells settle down at the bottom because of their dense coacervate core. Note that the vesicle layer is present in both samples, but the pellet is present only in the hybrid protocell sample, which can be visualized in the zoomed image. Free amphiphiles also remain in the supernatant because of their small size.

**Supplementary Table 1.** Masses corresponding to different single-chain amphiphiles that were detected during the LC-MS analysis of heterogeneous hybrid protocell membranes.

| Reaction | Chemical species | Exact mass | Ion type | Theoretical mass | Observed mass | Error (ppm) |
|---|---|---|---|---|---|---|
| OA + GMO hybrid protocells | OA | 282.2559 | $[M-H]^-$ | 281.2486 | 281.2480 | - 2.1 |
| | GMO | 356.2927 | $[M+HCOO]^-$ | 401.2909 | 401.2898 | - 2.7 |
| OA + MA hybrid protocells | OA | 282.2559 | $[M-H]^-$ | 281.2486 | 281.2462 | - 8.5 |
| | MA | 226.1933 | | 255.1860 | 225.1853 | - 2.7 |

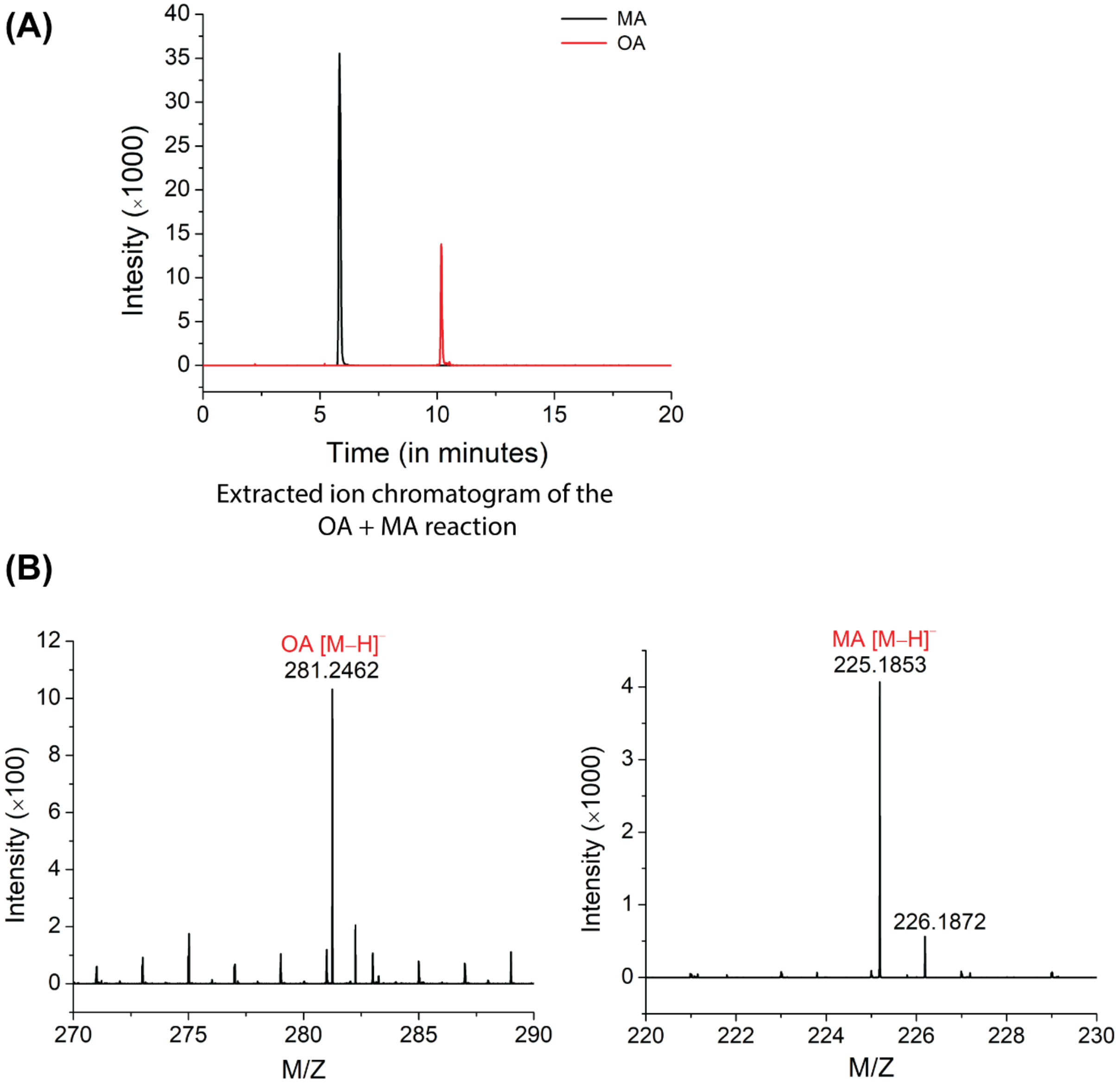


**Figure S6. Confirming the membrane heterogeneity of OA + MA hybrid protocells by mass spectrometry. (A)** The extracted ion chromatogram shows the peaks corresponding to MA and OA ions with retention times of ≈ 6 min and ≈ 10 min, respectively. **(B)** The corresponding mass spectra in negative ion mode show the expected masses for MA (225.1853) and its $^{13}C$ isotope (226.1872) (right spectrum) and OA (281.2462) (left spectrum).

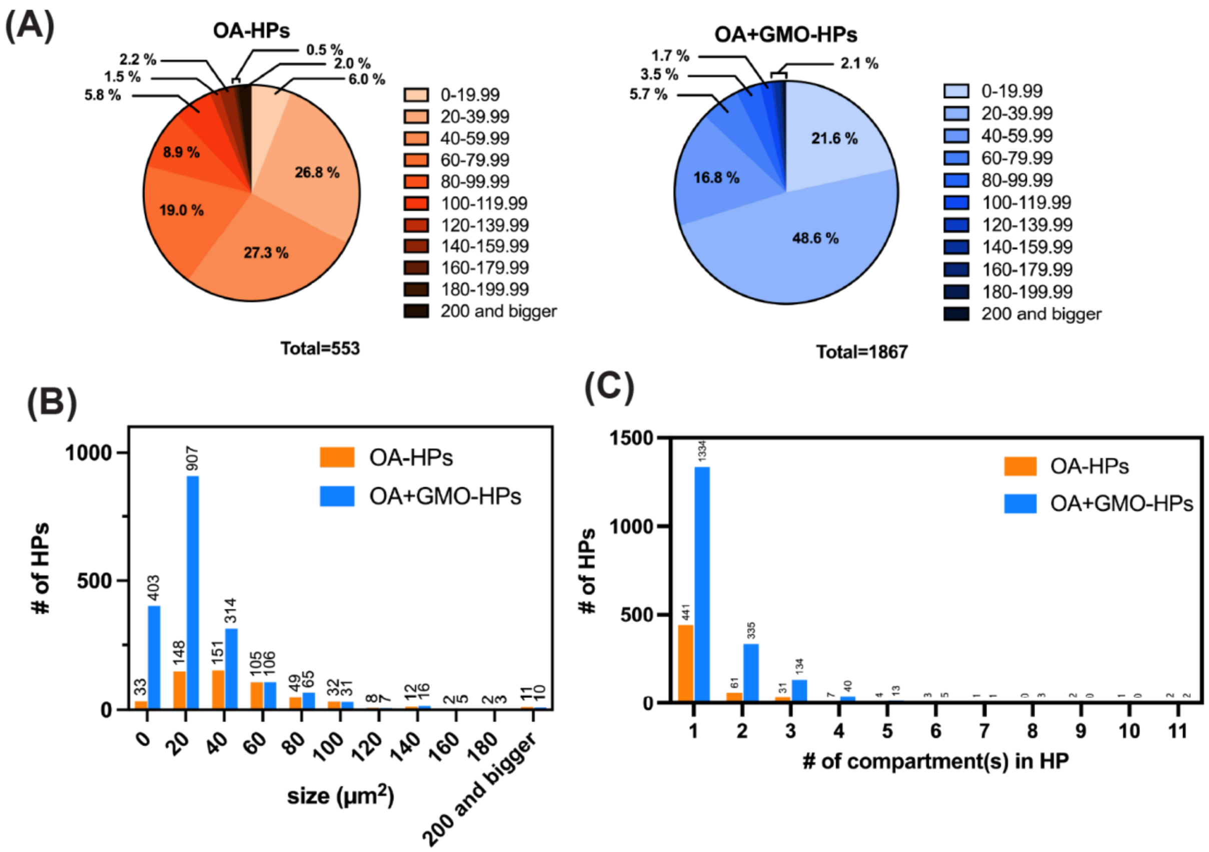


**Figure S7. Comparing the size and multicompartment formation efficiency of homogeneous and heterogeneous hybrid protocells. (A)** Pie charts depicting the size distribution of OA and OA + GMO HPs and the percentage of HPs observed within the specific size range. **(B)** Histogram showing size comparison of OA and OA + GMO HPs. **(C)** Histogram showing the comparison of OA and OA + GMO HPs in terms of the number of HPs with different counts of sub-compartments.

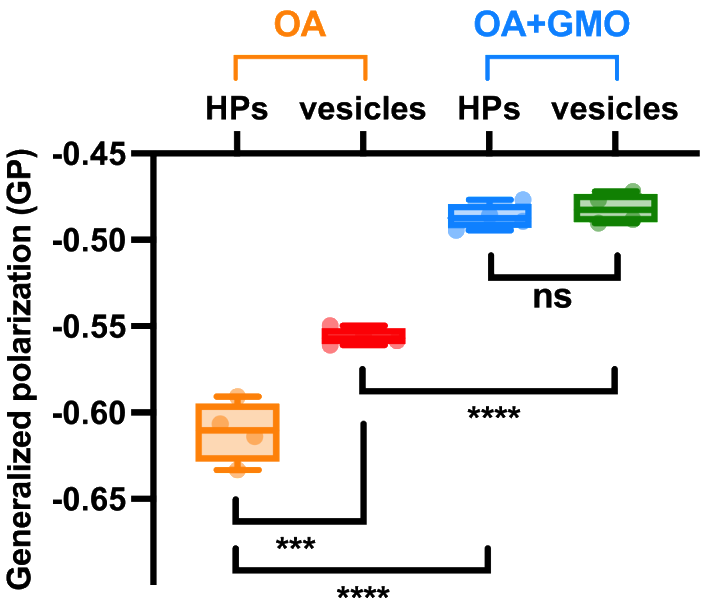


**Figure S8. Comparing the membrane order of hybrid protocells and vesicles.** Generalized polarization (GP) values of OA (9 mM) and OA + GMO (9 mM; 2:1 ratio) hybrid protocells and vesicles based on their Laurdan emission spectra. Data is represented as a box and whisker plot, where data points are shown alongside the box with center lines showing the medians, box limits indicating the 25th and 75th percentiles, and whiskers extending to the most extreme data points. Error bars represent standard deviation, and the statistical significance is calculated using a two-tailed unpaired t-test, where $n = 4$, *** represents $p < 0.001$, **** represents $p < 0.0001$, and ns means not significant.

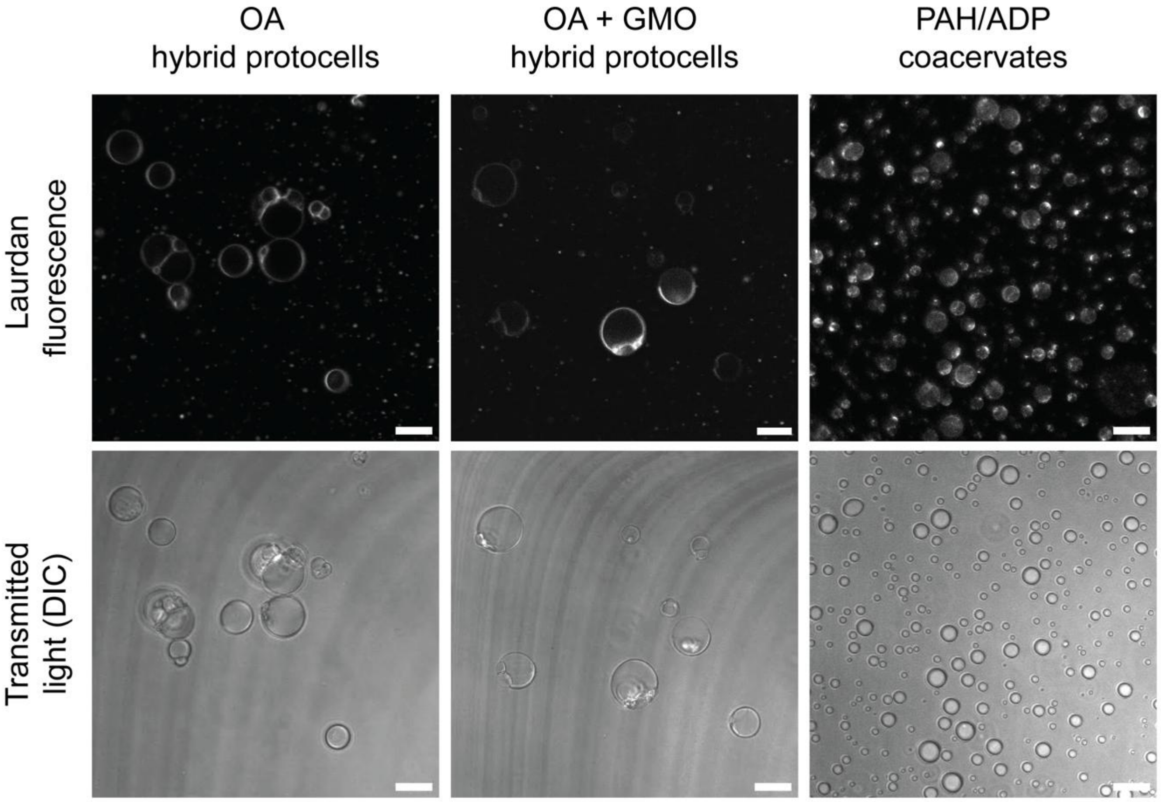


**Figure S9. Localization of Laurdan dye in hybrid protocells and coacervates.** Laurdan dye was added to OA (9 mM) and OA + GMO (9 mM; 2:1 ratio) hybrid protocells and also to the PAH/ADP coacervates (40 mM total charge; 1:1 charge ratio). The concentration of Laurdan dye was the same as that used for estimating the membrane order (9 µM; 1:1000 ratio with total lipid concentration). Fluorescence images were collected with Ex = 405 nm and Em = 450 nm to 550 nm. The scale bar is 10 µm.

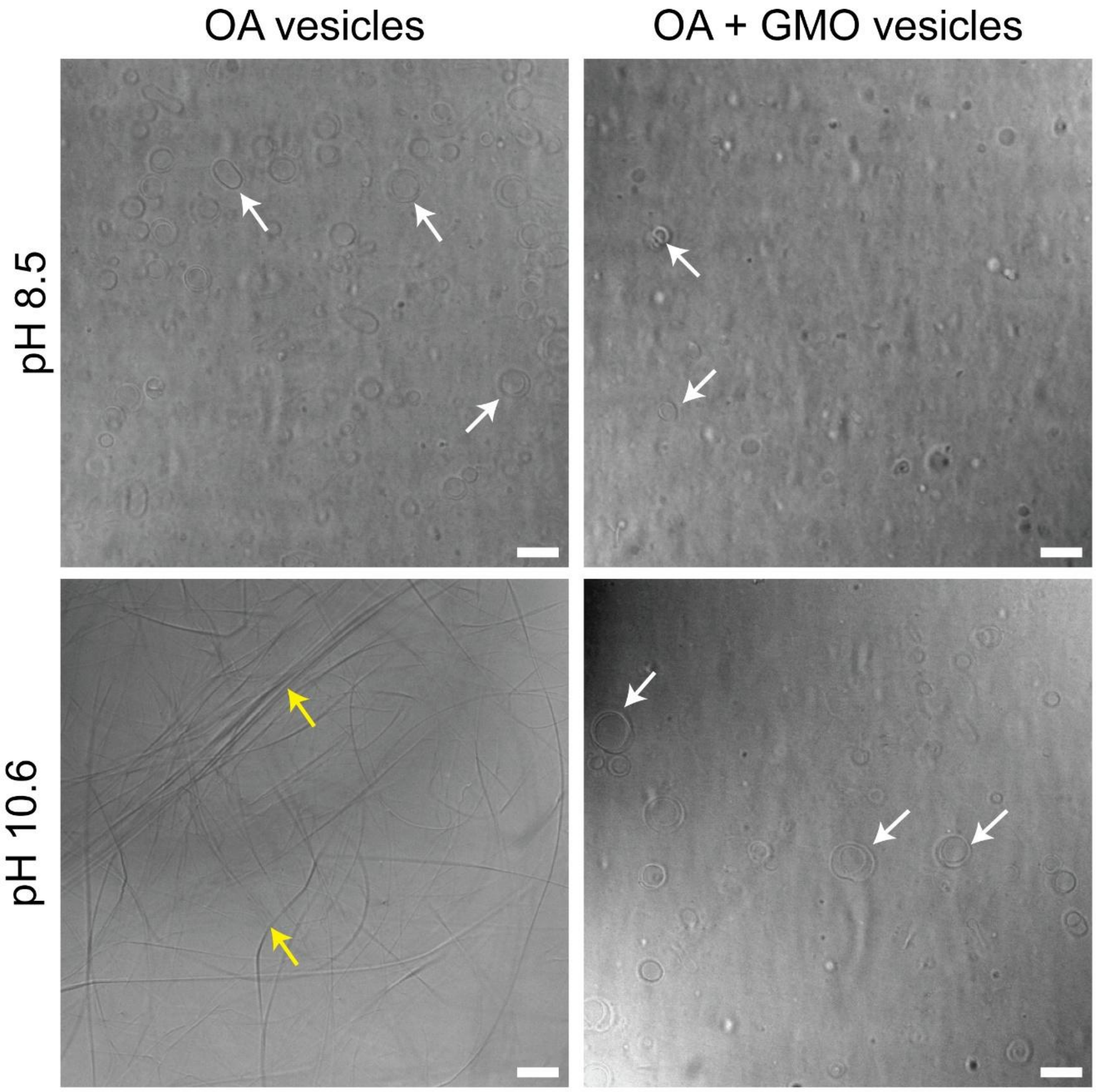


**Figure S10. Alkaline pH stability of OA and OA + GMO vesicles.** OA (9 mM) and OA + GMO (9 mM; 2:1 ratio) vesicles were prepared in 100 mM bicine buffer, pH 8.5. Images were acquired using a differential interference contrast (DIC) channel. Examples of vesicles and crystals are indicated by white and yellow arrows, respectively. The scale bar is 10 μm.

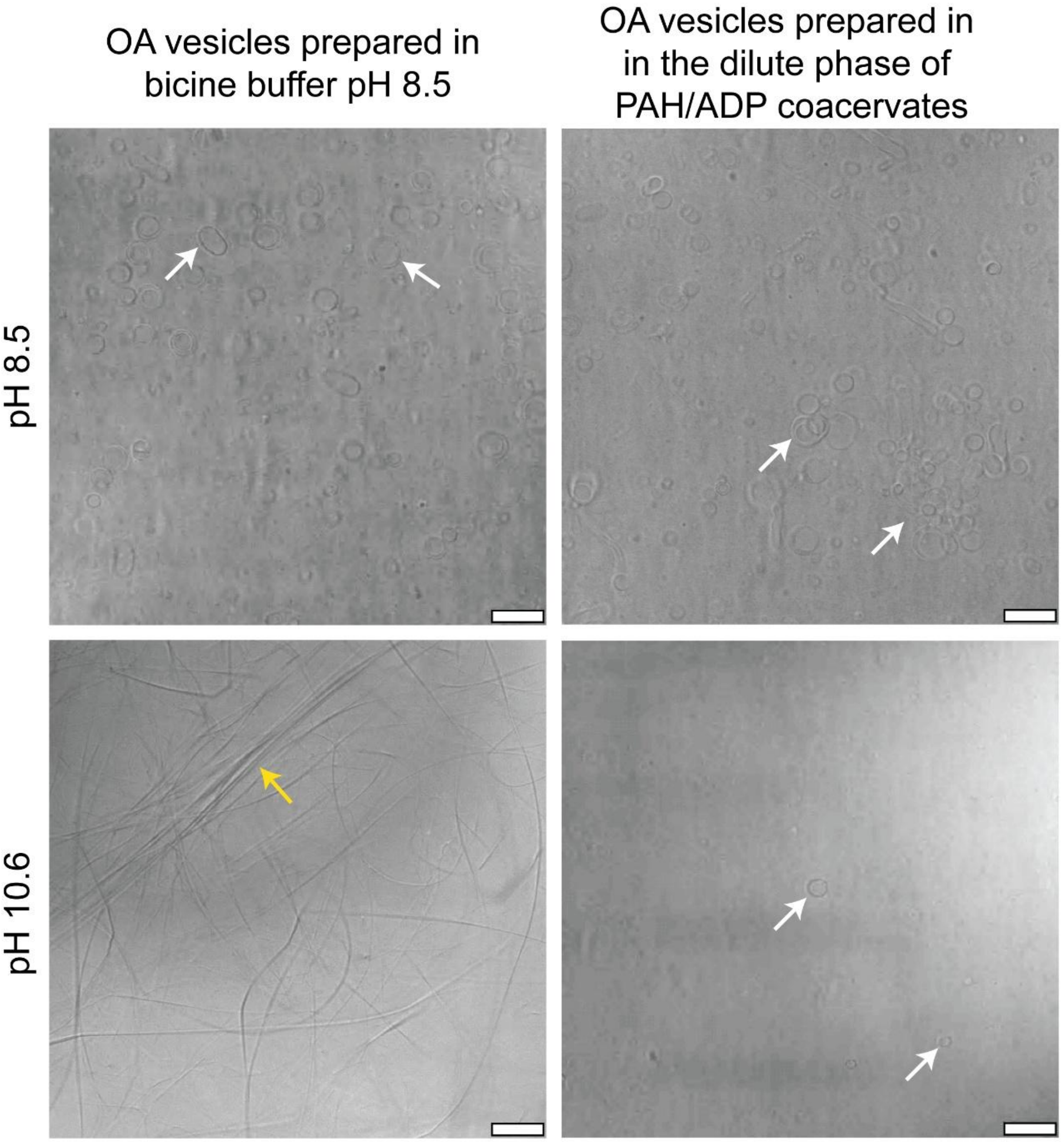


**Figure S11. Comparing the alkaline pH stability of OA vesicles prepared in bicine buffer versus the dilute phase of PAH/ADP coacervates.** OA vesicles (9 mM) were prepared in 100 mM bicine buffer alone, and the dilute phase of PAH/ADP coacervates (40 mM total charge; 1:1 charge ratio), prepared in the same bicine buffer. Both solutions had a starting pH of 8.5. Images were acquired using a differential interference contrast (DIC) channel. Examples of vesicles and crystals are indicated by white and yellow arrows, respectively. The scale bar is 10 μm.

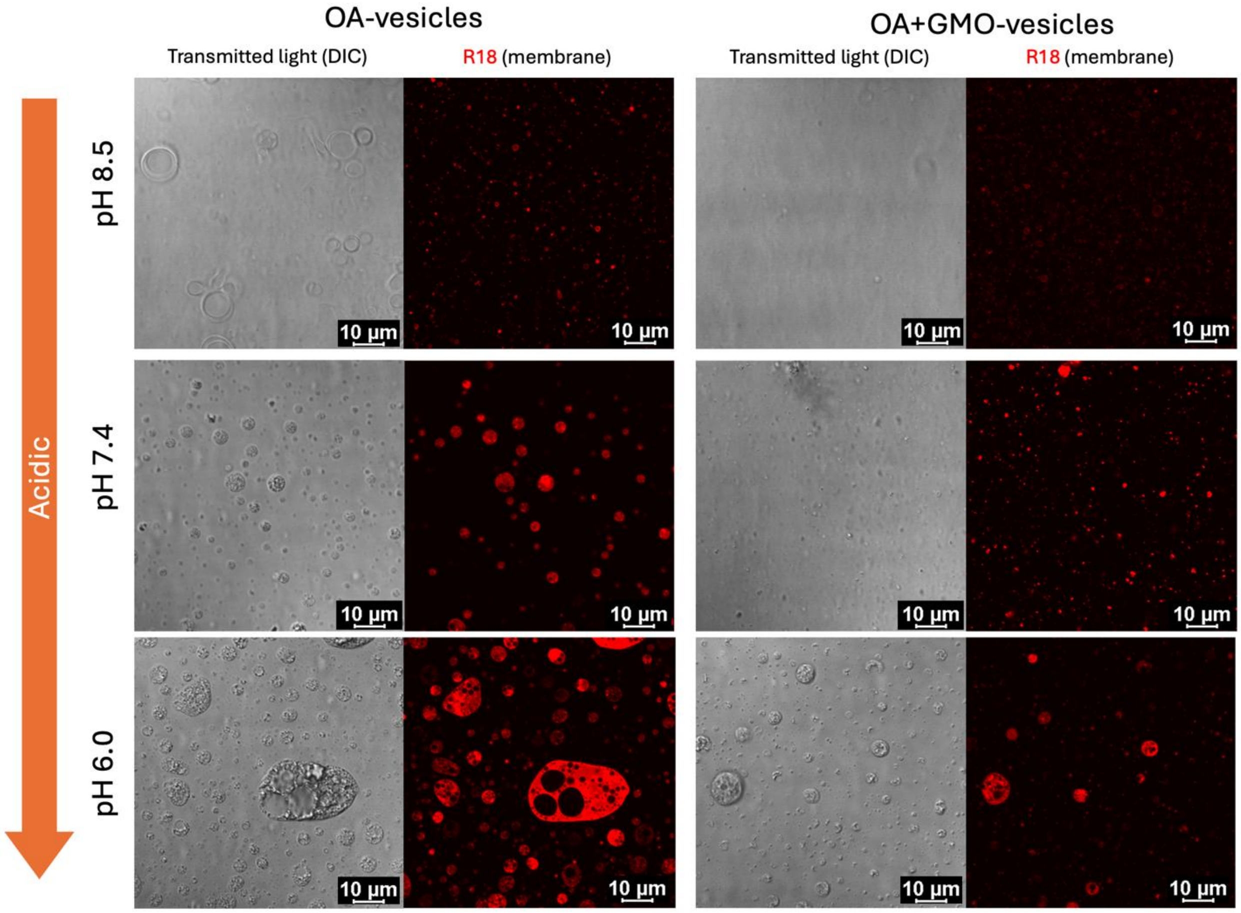


**Figure S12. Acidic pH stability of OA and OA + GMO vesicles.** OA (9 mM) and OA + GMO (9 mM; 2:1 ratio) vesicles were prepared in 100 mM bicine buffer, pH 8.5. Images were acquired using a differential interference contrast (DIC) channel and R18 fluorescence channel (indicating membrane). Both OA and OA + GMO vesicles exhibited a clear transition into oil droplets as the pH decreased. This morphological instability was more significant in the homogeneous OA vesicles compared to the heterogeneous OA + GMO vesicles.

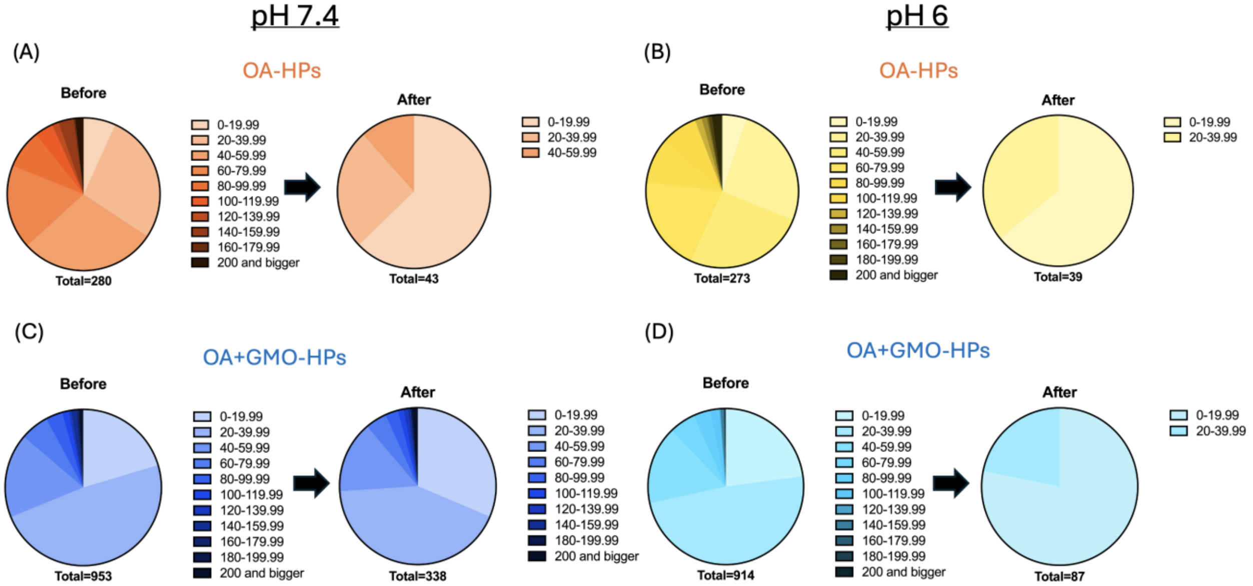


**Figure S13. Shift in size distribution and survival of protocells upon acidification.** The pH of OA HPs **(A, B)** and OA+GMO HPs **(C, D)** was decreased from 8.5 to 7.4 and 6.0, respectively. OA+GMO HPs demonstrated significantly higher survival rates (338 at pH 7.4; 87 at pH 6.0) compared to pure OA HPs (43 at pH 7.4; 39 at pH 6.0) at reduced pHs. While OA+GMO HPs maintained greater size heterogeneity at pH 7.4 compared to pure OA membranes, both systems exhibited similar levels of heterogeneity when the pH was lowered to 6.0.

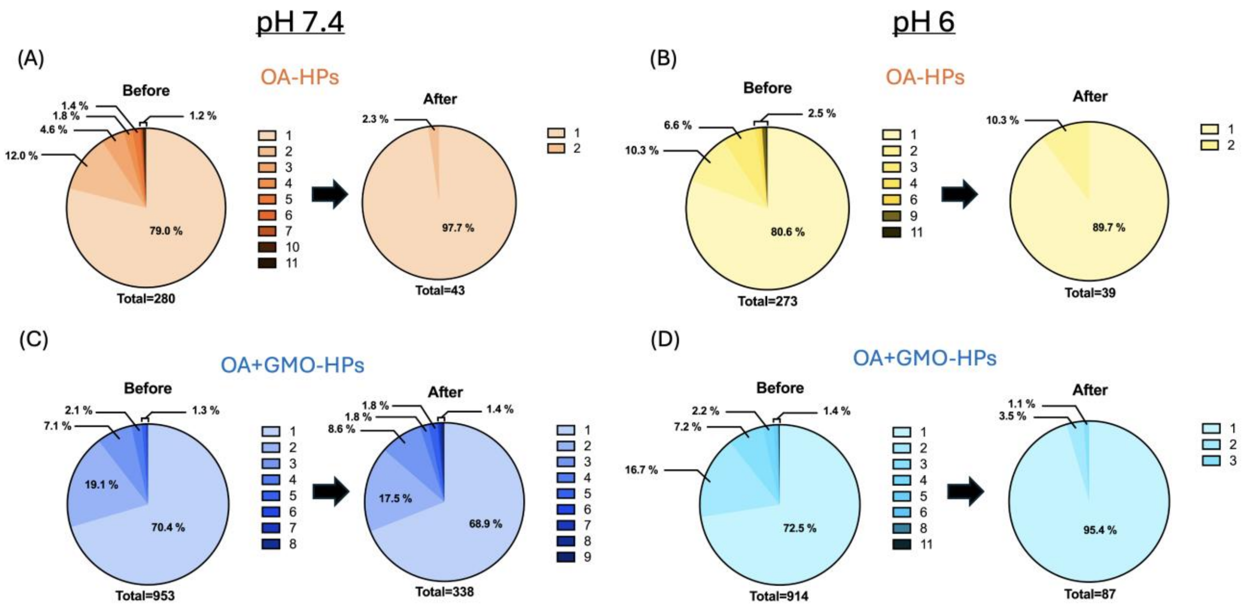


**Figure S14. Shift in internal compartment distribution and survival of protocells upon acidification.** The pH of OA HPs **(A, B)** and OA+GMO HPs **(C, D)** was decreased from 8.5 to 7.4 and 6.0, respectively. OA + GMO HPs retained higher compartment heterogeneity across both tested pH values compared to OA-HPs, which transitioned toward more simplified internal structures.
